# Human POLD3 coordinates leading and lagging strand in Mitotic DNA Synthesis

**DOI:** 10.64898/2025.12.15.694317

**Authors:** Demetrio Turati, Vasilis S. Dionellis, Laurence Tropia, Thanos D. Halazonetis

**Affiliations:** Department of Molecular and Cellular Biology, University of Geneva, Geneva, Switzerland; Department of Visceral Surgery and Medicine, Inselspital and Department for BioMedical Research, University of Bern, Bern, Switzerland

## Abstract

Deregulation of origin firing and licensing, shortage of deoxyribonucleotides in the cell and interference between transcription and replication are among the causes of DNA replication stress in cancer cells. This leads in various ways (notably, through common fragile sites expression and incomplete replication of late-replicating genomic regions during S phase) to the necessity of finishing DNA replication in mitosis by a mechanism called Mitotic DNA synthesis (MiDAS), which is related to Break-Induced Replication (BIR). Even if it is of primary importance for cancer cells, the molecular mechanism of MiDAS, and generally of BIR, is not yet well understood. Recently, the third subunit of the eukaryotic DNA polymerase delta (POLD3) has been recognized as a key player of BIR, though its role is not yet clear. In this work, using a protocol established in our group to map at high resolution newly replicated DNA at MiDAS sites, we provide new insights into the molecular role of POLD3 in this repair mechanism. In particular, by analyzing MiDAS in mutant HeLa clones lacking the PCNA-interacting domain of POLD3, we demonstrate that the interaction between POLD3 and PCNA is required for coordinating leading and lagging strand synthesis in MiDAS. This work represents an important step forward towards the comprehension of the molecular mechanisms of Mitotic DNA synthesis, the full understanding of which is of primary importance for the possible development of novel cancer therapies targeting BIR-related pathways.

## INTRODUCTION

POLD3 is the third subunit of the DNA polymerase δ complex, and is commonly considered responsible for lagging strand synthesis during S-phase DNA replication in eukaryotes ^1^. Human, POLD3 (also known as p66 due to its apparent molecular weight in SDS-PAGE ^2^) is a 466-residues protein ^3^ with a molecular weight of 51.4 kDa. It encompasses, from the N- to the C-terminus, a POLD2-interaction domain (residues 1-144 ^4^), which is also the only structured part of the protein, a Rev1-interaction motif (RIR, residues 231-246 ^5^), a Protein Phosphatase 1 (PP1)-interaction motif (residues 302-306 ^6^), a DNA polymerase α-interaction motif (DPIM, residues 394-402 ^7^) and a PCNA-interaction motif (PIP-box, residues 456-463 ^8^).

Together with POLD4 (p12), POLD3 has been commonly referred to as a non-essential subunit of the mammalian DNA polymerase δ holoenzyme, since the originally purified heterodimer composed of the POLD1(p125) and POLD2(p50) subunits is able to perform DNA synthesis in vitro by itself ^9^. However, subsequent investigation showed that the PCNA-dependent stimulation of DNA polymerase δ activity in vitro strongly relies on the presence of POLD3 ^10^ and that POLD3 knock-out is lethal in vivo ^11^. This indicates that the third subunit actually plays an essential role in the DNA polymerase δ holoenzyme function in the cellular context, so that p66 should be better considered a *sub-*essential subunit of the complex rather than a non-essential one. Interestingly, previous work of our group ^12,13^ showed that depletion of p66 increases the levels of DNA damage and impairs S-phase entry and progression in cells with DNA replication stress but not in healthy cells, suggesting that POLD3 could play an additional role in Break-Induced Replication (BIR) that is not required in normal DNA replication. Indeed, POLD3 depletion negatively affects the activity of BIR ^12,13^ and BIR-related mechanisms such as break-induced telomere synthesis ^14^ and Mitotic DNA Synthesis (MiDAS) ^15,16,17^, a recently discovered RAD52-dependent BIR-like pathway involved in DNA repair at Common Fragile Sites (CFS) and at Under-replicated DNA Regions (UDRs) during mitosis ^16,17^.

The discovery of MiDAS represents an important step forward in the study of BIR-related pathways. Indeed, we now know that, besides being involved in the repair of single-ended double strand breaks rising from transcription-replication collisions at oncogene-induced replication origins ^12,13,18^ and in the elongation of telomeres in telomerase-negative tumors through the Alternative Lengthening of Telomeres (ALT) pathway ^19^, BIR-like mechanisms are also involved in another mechanism on which cancer cells with DNA replication stress strongly rely on ^20^. This further increases the interest in BIR-related pathways as targets for novel cancer therapies, and triggers a strong need for a better understanding of this class of repair mechanisms.

The identification of MiDAS also provides a new powerful technical tool to study BIR-like pathways. Break-Induced Replication at oncogene-induced origins cannot be easily studied due to the concomitant presence of normal S-phase DNA synthesis, which represents an important source of background. As far as ALT is concerned, this BIR-like pathway happens at highly repeated sequences, the telomeres, which makes difficult to unequivocally identify the newly replicated DNA. Studying BIR at MiDAS sites allows to overcome both these problems: MiDAS happens in mitosis, thus representing the only source of novel DNA synthesis in the cells, and mainly at large genes ^21^, which are unique genomic sequences. Following this idea, a method for studying MiDAS at a sequencing level has been established in our group ^21,22^ based on a protocol by Hickson and colleagues that was used to study MiDAS by fluorescence microscopy ^16,17^. By using this new method we were able to provide mechanistic insights for DNA replication during MiDAS, and to map MiDAS sites on the genome at high resolution in U2OS, HeLa and HS68 cell lines. We were able to observe that, in a cell population, MiDAS starts on both sides of the CFS or UDR, and that the two forks approach each other towards the center of the under-replicated region. Moreover, we have also noticed that in HeLa cells, but not in U2OS or HS68 cells, DNA synthesis during MiDAS has some delay in lagging strand synthesis or Okazaki fragments ligation, which is observable as a directionality in the sequencing reads ^21^.

Although the involvement of POLD3 in BIR-related pathways is clear, its precise molecular role in these mechanisms remains elusive, and could even be different in the various BIR subtypes. Possibly, POLD3 could be involved in mediating the recruitment of various BIR players at the repair site. Recent studies suggest that MiDAS might be initiated by the REV1-Polζ translesion synthesis complex, and that Polδ might be recruited to the MiDAS site afterwards through the interaction between POLD3 and REV1 and between POLD2 and REV7 ^15^. However, this might not be the case for ALT, since previous data showed that REV1 and REV3 are not needed for break-induced telomere synthesis ^14^. The same work suggested that recruitment of Polδ at telomeres could be mediated by the interaction between POLD3 and PCNA, since deletion of the PCNA-interacting motif of POLD3 impairs the recruitment of the whole DNA polymerase δ complex to telomeres after TRF-FokI-induced damage. Interestingly, this study also suggested that DNA polymerase δ synthesizes both leading and lagging strand during break-induced telomere synthesis, in agreement with previous work showing that Polδ is able to synthesize both leading and lagging strands in BIR in budding yeast ^23^.

Conscious of the potential of the protocol we set up to investigate DNA replication at MiDAS sites ^21,22^, we decided to use it to better understand the role of POLD3 in BIR. First, we generated several HeLa cell lines with mutations in various functional motifs of p66. Then, we studied MiDAS in the mutant and wild type cell lines. Notably, we observed that, upon deletion of the PIP-box of POLD3, MiDAS is not abolished, differently from what has been observed for break-induced telomere synthesis by Dilley and colleagues ^14^. However, we found that this mutation heavily affects the coordination between leading and lagging strand synthesis in MiDAS, although S-phase DNA synthesis is not affected in this sense. This observation sheds new light on the involvement of POLD3 in MiDAS, suggesting that p66 might have different functions in BIR and in normal DNA replication.

## RESULTS

### Generation of HeLa cell lines lacking functional motifs of POLD3

To understand the role of POLD3 in MiDAS, we generated by CRISPR-Cas9 technology five HeLa clones lacking various functional motifs of POLD3 (Figures 1a, Supplementary Figure 1 and Supplementary Table 1): two clones lacking the PCNA-interacting motif (clones 23 and 2C4), two clones lacking the PP1 interaction motif (clones 2E4 and 3H2) and one clone lacking the RIR motif (clone 3D1). Even if the mutations slightly affected the expression of POLD3 in some of the clones, the overall expression levels of the mutant proteins were comparable to the wild type (Supplementary Figure 2). An attempt to produce a POLD3 KO HeLa clone failed due to the lethality of the mutation (also observed by Dilley and colleagues ^14^), consistent with POLD3 being essential for normal DNA replication ^11^.

**Figure 1:**
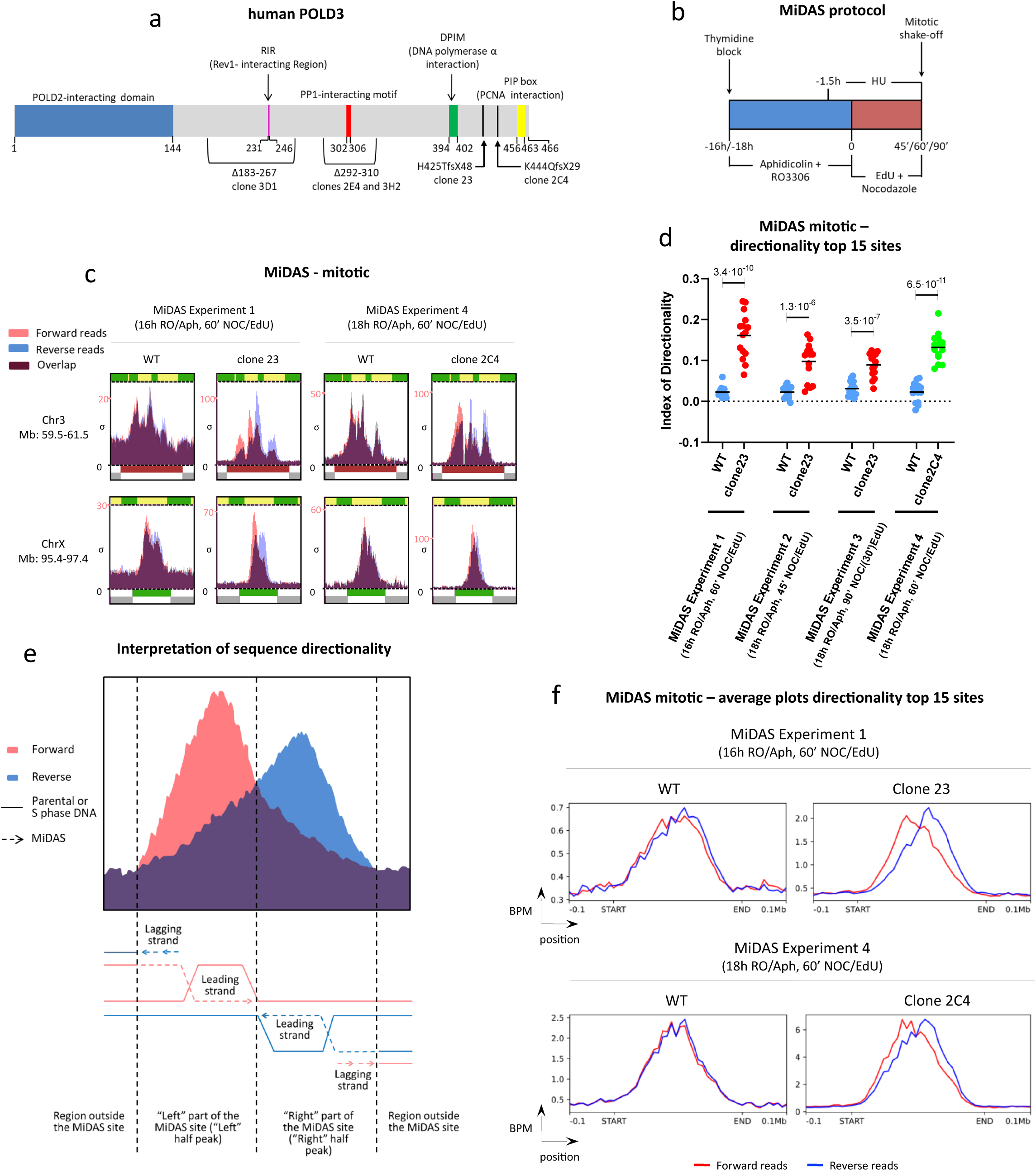
Sequence directionality at MiDAS sites upon deletion of the PCNA-interacting motif of POLD3. **a)** Structure of human POLD3 with indication of its functional domains and motifs, and of the mu-tations of the various mutant clones we produced. **b)** Schematic representation of the adapted MiDAS protocol used in this work. **c)** Results of MiDAS experiments performed on HeLa POLD3^delPIP-box^ mitotic cells, clone 23 and clone 2C4 versus wild type. Peaks from forward and reverse reads are shown in red and blue, re-spectively. **d)** Dot plot showing the Index of directionality of the top 15 MiDAS sites (each dot is a MiDAS peak) for the HeLa POLD3^delPIP-box^ mutant clones and wild type in each of the MiDAS experiments. The p-values reported on the significance bracket come from a Student’s t-test performed between each mutant clone and the wild type control, considering each MiDAS peak as a repeat (statistical significance: p-value<0.05). The Index of Directionality (IoD) for each peak can range between −1 and 1 and is calculated as:

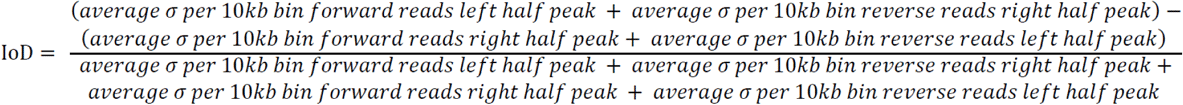 An IoD greater than 0 indicates positive directionality (excess of forward reads in the left half peak and/or excess of reverse reads in the right half peak), an IoD smaller than 0 indicates negative direc-tionality (excess of forward reads in the right half peak and/or excess of reverse reads on the left half peak) and IoD equal to 0 indicates no directionality (equal number of forward and reverse reads in both left and right half peaks). For each experiment, a horizontal bar indicates the average IoD value of each cell line. **e)** Proposed mechanism to explain the directionality observed at MiDAS sites after mutation of the PCNA-interacting motif of POLD3. The delayed production and/or ligation of Okazaki Fragments at MiDAS forks leaves nicks in the lagging strand, making not possible to sequence fragments com-ing from this strand with Next Generation Sequencing. This causes an excess of sequencing reads mapping on the leading strand of the MiDAS forks. **f)** Average plots of the top 15 peaks for clone 23 (experiment 1), clone 2C4 (experiment 4) and wild type, covering the MiDAS region (from the list provided in ^21^) ±0.1 Mb. The start and end points of the MiDAS peaks were aligned to standardize the different genomic size of the various MiDAS re-gions. Average peaks coming from forward and reverse reads are colored in red and blue, respec-tively. BPM: bins per million mapped reads. BPM (per bin) = number of reads per bin / sum of all reads per bin (in millions).

### POLD3 mutations do not impair MiDAS activity

To understand the possible role of the different POLD3 motifs in BIR we performed a MiDAS experiment as in ^21,22^ in all the HeLa POLD3^mut^ cell lines we generated, as well as in the parental HeLa cells (Figure 1b). Briefly, cells were blocked in G1/S by thymidine block and subsequently released in S-phase in the presence of aphidicolin and the Cdk1 inhibitor RO3306 for either 16 or 18 hours. Aphidicolin, an inhibitor of B-family DNA polymerases, induces DNA replication stress, CFS expression ^24^ and MiDAS ^17^, while RO3306 blocks cells at the G2/M boundary. After washing away the drugs, cells were released into mitosis in the presence of hydroxyurea (HU), 5-Ethynyl-2’-deoxyuridine (EdU) and nocodazole for 45, 60 or 90 minutes (in the latter case, EdU was added only for the last 30 minutes). HU blocks S-phase DNA synthesis in non-mitotic cells that might be contaminating the mitotic cells, reducing the background signal of EdU incorporation; EdU, in turn, is used to label newly synthesized DNA in mitosis; whereas, nocodazole arrests cells in mitosis ^17^. Mitotic and non-mitotic (attached) cells were collected by mitotic shake-off and trypsinization respectively, and an EdU-seq protocol was performed on the two cell populations as described in ^18^. The EdU-labelled DNA, corresponding to newly replicated DNA in mitosis, was sequenced by single-end Next Generation Sequencing, and the reads obtained were aligned on the human reference genome. The list of MiDAS sites for HeLa cells reported in ^21^ was used as a reference to analyze the MiDAS peaks in the sequencing alignment.

A first analysis of the sequencing results aimed to check the possible impairment of MiDAS upon deletion of the functional domains of POLD3. Surprisingly, DNA replication peaks at MiDAS sites were detected in all the clones, meaning that none of the mutations had a strong negative effect on Mitotic DNA Synthesis (data not shown). This is in contrast with previous work on break-induced telomere synthesis showing that deletion of the PIP-box of POLD3 prevents its recruitment to TRF1-FokI-induced telomere damage ^14^, and that mutation of the RIR motif of POLD3 impairs its recruitment to chromatin at MiDAS sites and reduces MiDAS activity ^15^. These different observations might be explained in the light of the fact that the first above-cited study investigated the role of POLD3 in break-induced telomere synthesis, instead of MiDAS, whereas the second study studied U2OS, rather than HeLa cells.

Even if MiDAS was present in all the cell lines, we observed some differences in the height of the peaks between the parental cells and mutant clones (data not shown). However, since we did not observe consistency among the various experiments and even among different MiDAS sites in the same experiment, we assumed that these differences were most probably related to technical variability.

### Deletion of the PCNA-interacting motif of POLD3 causes sequence directionality at MiDAS sites

A previous work from our group ^21^ showed that in HeLa cells, but not in U2OS or HS68, MiDAS peaks are characterized by directionality of sequencing reads, which means, there is an excess of sequencing reads mapping on the forward strand of the genome on the left part of the MiDAS peak, and vice versa on the right part. This might be due to some delay in Okazaki fragment synthesis or ligation in HeLa cells, which makes the lagging strand not amplifiable during the library preparation for single-end next generation sequencing (Figure 1e, Supplementary Figure 3). Performing this kind of analysis in the POLD3 mutant clones we observed that both clones lacking the PCNA interaction domain of POLD3 (clone 23 and clone 2C4), but not the other mutant clones, had a much more pronounced sequence-read directionality than the parental cells, as indicated by a bigger shift between the forward and reverse peaks at each MiDAS site (Figures 1c and 2a, Supplementary Figure 4).

**Figure 2:**
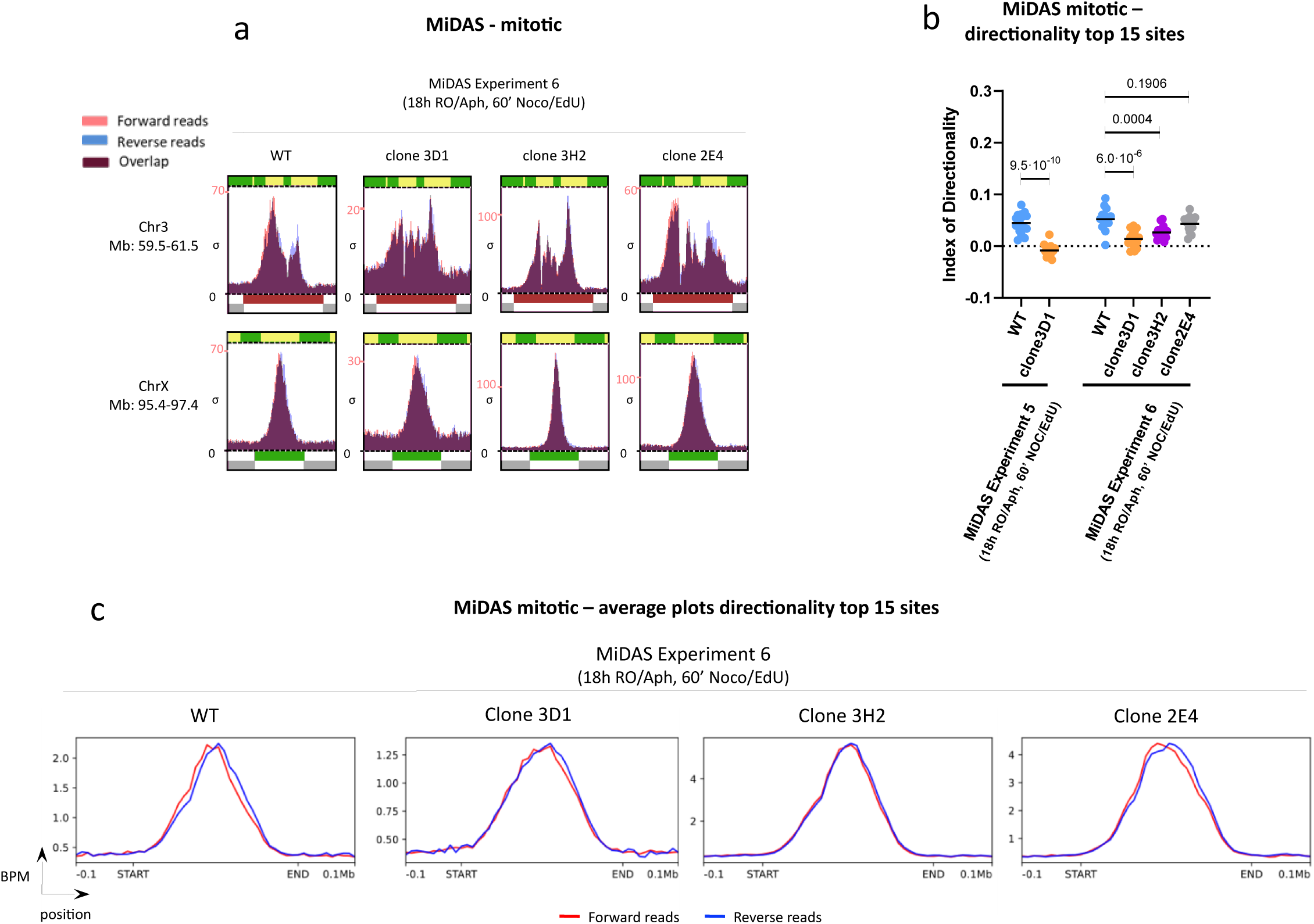
Analysis of sequence directionality at MiDAS sites upon deletion of the RIR-motif or the PP1-interacting motif of POLD3. **a)** Results of MiDAS experiments performed on HeLa POLD3^delRIR^ mitotic cells (clone 3D1) and HeLa POLD3^delPP1^ mitotic cells (clones 3H2 and 2E4) versus wild type. Peaks from forward and re-verse reads are shown in red and blue, respectively. **b)** Dot plot showing the Index of directionality of the top 15 MiDAS sites (each dot is a MiDAS peak) for the mutant clones and wild type ineach of the MiDAS experiments. The p-values reported on the significance bracket come from a Student’s t-test performed between each mutant clone and the wild type control, considering each MiDAS peak as a repeat (statistical significance: p-value<0.05). For each experiment, a horizontal bar indicates the average IoD value of each cell line. **c)** Average plots of the top 15 peaks for clones 3D1, 3H2, 2E4 and wild type, covering the MiDAS region (from the list provided in ^21^) ±0.1 Mb. The start and end points of the MiDAS peaks were aligned to standardize the different genomic size of the various MiDAS regions. Average peaks coming from forward and reverse reads are colored in red and blue, respectively. BPM: bins per million mapped reads. BPM (per bin) = number of reads per bin / sum of all reads per bin (in mil-lions).

To quantify the directionality and to make an objective comparison between the parental cells and mutant clones we calculated an Index of Directionality (IoD) for each MiDAS peak (see Materials and Methods). The IoD can be equal to 0 if no directionality is present, or can assume a positive or negative value in case of positive (excess of forward reads on the left side and excess of reverse reads on the right side of the MiDAS peak) or negative (excess of reverse reads on the left side and an excess of forward reads on the right side of the MiDAS peak) directionality, respectively. We then compared the calculated IoDs of the MiDAS peaks of every clone with the ones of the parental cells. Since not all the MiDAS peaks reported for HeLa cells in ^21^ were present in our datasets, we compared the IoDs of a subset of 15 MiDAS peaks for each cell line and for each experiment, selected on the basis of the intensity of the signal and of the signal/background ratio (see Extended Data). As was already evident by simple inspection of the MiDAS peaks, comparison of the IoDs confirmed the presence of a statistically significant higher level of positive directionality in the mutant clones missing the PCNA-interacting motif of POLD3 with respect to the parental cells (Figure 1d, Supplementary Table 2). Surprisingly, clone 3D1, which misses the RIR motif of POLD3, and clone 3H2, which is one of the two clones missing the PP1-interacting motif, showed significant lower IoDs than the parental cells. On the contrary clone 2E4, namely the second clone with a deletion covering the PP1-interacting motif, did not show any significant difference in terms of IoD with respect to the parental cells (Figure 2b). To confirm these observations avoiding the possible biases due to the selection of the 15 best peaks, we repeated this analysis using the whole set of 206 MiDAS sites identified in HeLa cells in our previous study ^21^ (Supplementary Figure 7). This broader analysis confirmed the significance of the higher IoDs detected in clones 23 and 2C4 and the lower levels of directionality observed in clone 3D1, while the difference between the parental cells and clone 3H2 lost its significance. This suggests that the loss of the PP1-interacting motif of POLD3 does not actually affect directionality in MiDAS. Genome-wide average peaks of the most efficient 15 peaks (Figures 1f and 2c, Supplementary Figure 6) and of the whole set of MiDAS sites reported in HeLa cells (Supplementary Figure 8) further confirmed these observations.

Since the most striking and evident phenotype was due to the deletion of the PIP-box of POLD3 in clones 23 and 2C4, we decided to investigate more these two clones.

### The POLD3^delPIP-box^ mutants do not show directionality in S-phase DNA synthesis

To understand if the phenotype we observed in the HeLa POLD3^delPIP-box^ mutants was MiDAS-specific, or if the mutation was also affecting normal DNA replication, we also sequenced non-mitotic (attached) cells of clone 23 after having performed a MiDAS protocol. Newly replicated, EdU-labelled DNA in attached cells corresponds to late S-phase replication. Interestingly, no directionality was detected, showing that the deletion of the PIP-box of POLD3 has an effect on MiDAS, but not on S-phase DNA replication (Supplementary Figure 5).

To further probe directionality in S-phase cells, we blocked parental cells and mutant clones in prometaphase with nocodazole for 10 hours; then, collected the mitotic cells by mitotic shake-off and released them in G1 in the presence of HU for 13 hours, to block them at the beginning of S-phase. Next, the cells were released in S-phase for 45’, 90’ or 135’, the last 30’ of which were in the presence of EdU (Figure 3a). Pull down of the EdU-labelled DNA and single-end Next generation Sequencing was performed, and forward and reverse sequencing reads were mapped on the human reference genome to detect any directionality in S-phase replication forks. We focused the subsequent analysis on a selection of 15 constitutive, early S phase replication origins from the list provided for U2OS cells in ^18^ (see Extended Data), selected on the basis of their efficiency and of the absence of nearby origins, since overlapping replication forks moving in opposite directions could potentially hide directionality. As shown in Figure 3b and Supplementary Figure 9, no directionality was detected either in the parental cells or in the mutant clones. The IoDs also did not reveal any significant difference in directionality (Figure 3c).

**Figure 3:**
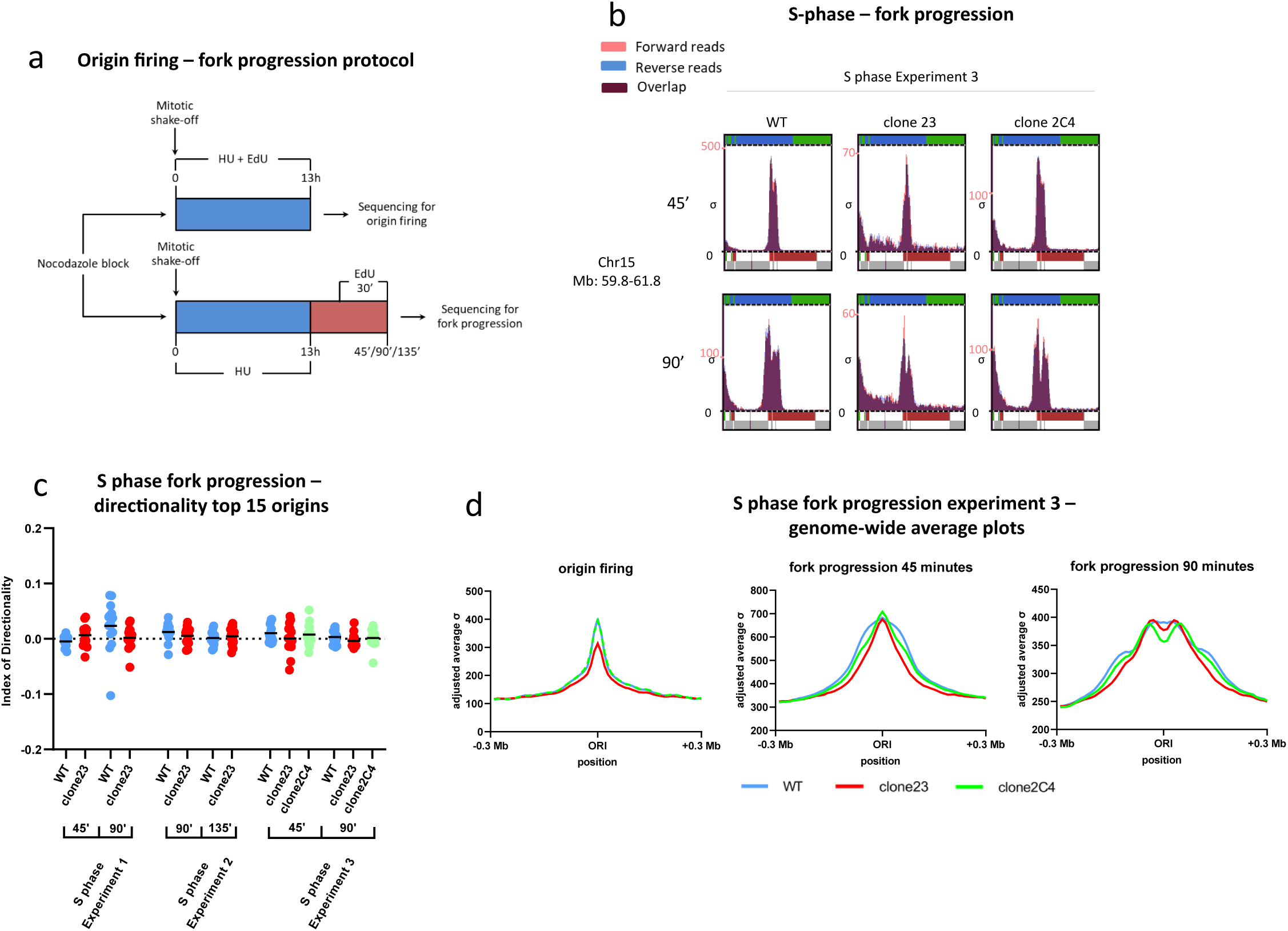
Deletion of the PCNA-interacting motif of POLD3 does not cause directionality in S-phase DNA synthesis. **a)** Schematic representation of the protocol used to investigate origin firing and fork progression in the mutant clones and wild type. **b)** Results of the fork progression experiments for a representative replication origin. Peaks coming from forward and reverse reads are colored in red and blue, respectively. **c)** Dot plot showing the Index of directionality of fork progression for the top 15 replication origins (each dot is an origin) of mutant clones and wild type in each of the fork progression experiments. **d)** Genome-wide average plots of the origin firing and fork progression experiments performed in the mutant clones versus wild type, on the basis of the replication origin list of U2OS cells reported in ^18^.

### The POLD3^delPIP-box^ mutation does not significantly affect S-phase DNA replication

The same data coming from the fork progression experiment, together with data about origin firing that we got from the same experiment (in the samples intended for origin firing analysis, EdU was added together with HU and they were collected at the end of the HU treatement), were used to check if the deletion of the PIP-box of POLD3 in the mutant clones was affecting the efficiency of origin firing and of fork progression. Average plots made using the whole list of replication origins for U2OS cells from ^18^, revealed a slightly lower efficiency in origin firing for clone 23 and a slightly slower fork progression for both mutant clones with respect to WT (Figure 3d, Supplementary Figure 10).

To see whether these slight differences in origin firing and fork progression had an effect on the cell cycle, we measured S-phase progression in the wild type and in the mutant clones by FACS. After thymidine block, cells were released in S-phase in presence of EdU for 3h, 6h, 9h or 12h (Figure 4a). EdU-PI staining was performed, and the FACS data produced (Figure 4b, Supplementary Figure 11) were analysed as described in Figure 4c to quantify S-phase progression at each time point. As shown in Figure 4d, although the mutant clones showed a slightly slower progression through S-phase, these differences were not statistically significant. Taken together, these data made us to conclude that deletion of POLD3 PIP-box does not affect S-phase DNA replication in a significant way.

**Figure 4:**
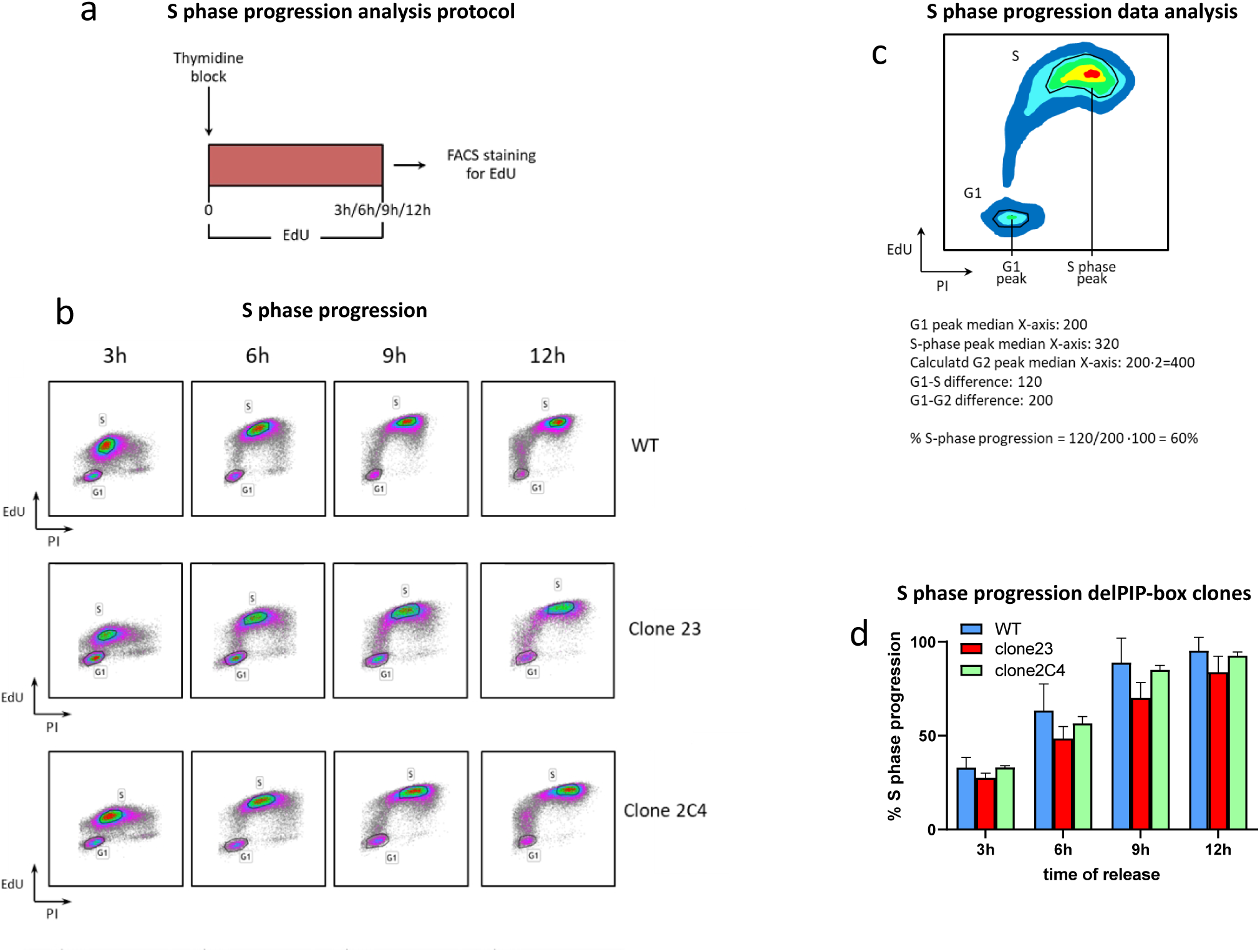
The POLD3^delPIP-box^ mutants show normal progression through S-phase. **a)** Schematic representation of the protocol used to investigate S-phase progression in the mutant clones and wild type by FACS. **b)** Results of the FACS experiments performed to investigate S-phase progression in the mutant clones and wild type. **c)** Method used to calculated the percentage of S-phase progression using the FACS data. To quan-tify S-phase progression, we calculated for each time point and for each cell line the distance cov-ered on the X axis (DNA content) by the center of the S-phase population with respect to the dis-tance between the G1 population and the calculated position of the G2 population (equal to double PI content of the G1 population). **d)** Histogram representing the percentage of S-phase progression of the mutant clones and wild type at various time points calculated on the basis of the FACS results.

## DISCUSSION

Previous work of our and other groups ^12–17^ showed that the POLD3 subunit of DNA polymerase δ is involved in BIR and BIR-related pathways. Differently from yeast Pol32, which deletion does not impair normal DNA replication ^26^, mammalian POLD3 seems to play a role also in S-phase, since we and others ^14^ were not able to produce POLD3 knock out cell lines, and knock out of POLD3 in vivo has heavy consequences on development and viability in mouse ^11^. However, POLD3 might play a different role in normal and in break-induced DNA replication. Indeed, depletion of POLD3 by siRNA in cultured U2OS cells triggers a strong increase in the levels of DNA damage with respect to control cells upon prolonged HU treatment or Cyclin E overexpression (i.e., upon induction of replication fork collapse) but not in absence of DNA replication stress ^12^. Moreover, POLD3 depletion impairs S-phase entry and replication fork progression only in cells overexpressing Cyclin E, while cells with normal expression of Cyclin E are not affected ^13^.

To better understand the specific role of POLD3 in BIR and BIR-related mechanisms, we investigated the effect of the deletion of various functional motifs of this protein on MiDAS. Specifically, we deleted by CRISPR-Cas9 technology the RIR motif, responsible for interaction of POLD3 with Rev1, the PP1-interacting motif and the PIP-box, which mediates interaction with PCNA. The clones we generated were analyzed for MiDAS as in ^21,22^.

Interestingly, none of the mutations abolished or significantly reduced MiDAS activity, in partial contrast with published data showing impairment of MiDAS or of break-induced telomere synthesis upon mutation of the RIR motif of POLD3 or deletion of the PIP-box ^14,15^. However, this does not necessarily represent a contradiction, considering that Wu and colleagues analyzed MiDAS in U2OS cells, which may have some differences with respect to MiDAS in HeLa ^15^, while Dilley and colleagues investigated a BIR-like mechanism different form MiDAS ^14^.

Notably, in the clones lacking the PCNA-interacting motif of POLD3 we detected a strong excess of sequencing reads mapping on the forward strand of the genome on the left side of each MiDAS peak and viceversa on the right side (positive directionality). Previous work of our group ^21^ observed that positive directionality is typical of MiDAS peaks in HeLa cells; however, the level of directionality we observed in the clones lacking the PIP-box of POLD3 is significantly higher than the WT. Due to the polarity of DNA synthesis, positive directionality means there is a surplus of reads coming from the leading strand of the replication fork with respect to those coming from the lagging strand on both sides of the MiDAS peaks, i.e., that leading and lagging strand synthesis are not coordinated. This can be due to a delayed Okazaki fragment synthesis or ligation; indeed, unligated Okazaki fragments cannot be amplified during the library preparation for next generation sequencing and thus cannot be sequenced.

Curiously, the levels of sequence directionality at MiDAS sites in clone 3D1 were significantly lower than in the WT. This might put our data in agreement with what reported by Wu and colleagues ^15^. Indeed, even if the analysis of the height of the MiDAS peaks did not show a reduced MiDAS frequency upon deletion of the RIR sequence (differently from what Wu and colleagues observed by EdU foci count upon mutation of the same motif ^15^), the decrease in sequence directionality might actually indicate a reduced rate or speed of leading strand synthesis. This might be due, as Wu and colleagues observed ^15^, to a reduced recruitment of POLD3 at MiDAS sites.

DNA polymerase δ is known to be responsible for lagging strand synthesis in normal DNA replication. Its activity is stimulated by the interaction of the holoenzyme with PCNA, which is mediated by POLD3 among the other subunits ^27,28^; thus, it makes sense that when the interaction between POLD3 and PCNA is compromised, the DNA polymerase δ holoenzyme activity at the replication fork is less efficient, causing a delay in lagging strand synthesis and/or Okazaki fragments ligation. However, even if the PCNA/DNA polymerase δ interaction is essential also for normal DNA replication, the deletion of the PIP-box of POLD3 only affects lagging strand synthesis in MiDAS, suggesting that the abolishment of the interaction between POLD3 and PCNA within the same Polδ complex might not be the explanation of the phenotype we observed. We might also exclude that the abolishment of the POLD3/PCNA interaction impedes the recruitment of the DNA polymerase δ at the MiDAS fork, since it has been suggested that recruitment of Polδ at MiDAS sites is mediated by interaction of POLD3 with Rev1 and of POLD2 with Rev3 of the Rev1-Polζ TLS complex ^15^. Also, if the recruitment of Polδ was abolished upon disruption of the POLD3/PCNA interaction, DNA synthesis would be probably impaired on both leading and lagging strand, since recent work suggest that Polδ synthesizes both leading and lagging strand in MiDAS^14^. Moreover, a loss of recruitment of the whole Polδ complex, even only on the lagging strand, would cause an accumulation of ssDNA that would at some point impair MiDAS activity, while in our experiments MiDAS still takes place.

Taking together our and other’s results, we propose a model (Figure 5) in which the POLD3 subunit of each Polδ complex interacts, through its unstructured C-terminal tail, with the PCNA trimer bound to the Polδ complex on the opposite strand of the MiDAS fork, providing the physical connection responsible for the coordination between leading and lagging strand synthesis in MiDAS. When the physical connection is lost, the two Polδ complex work independently from each other, and leading and lagging strand synthesis do not proceed coordinately anymore. Since in BIR the lagging strand is synthesized using the leading as template, it is conceivable that, upon loss of coordination between the two Polδ complexes, the discontinuous Okazaki fragment synthesis will proceed slower than the continuous synthesis of the leading strand.

**Figure 5:**
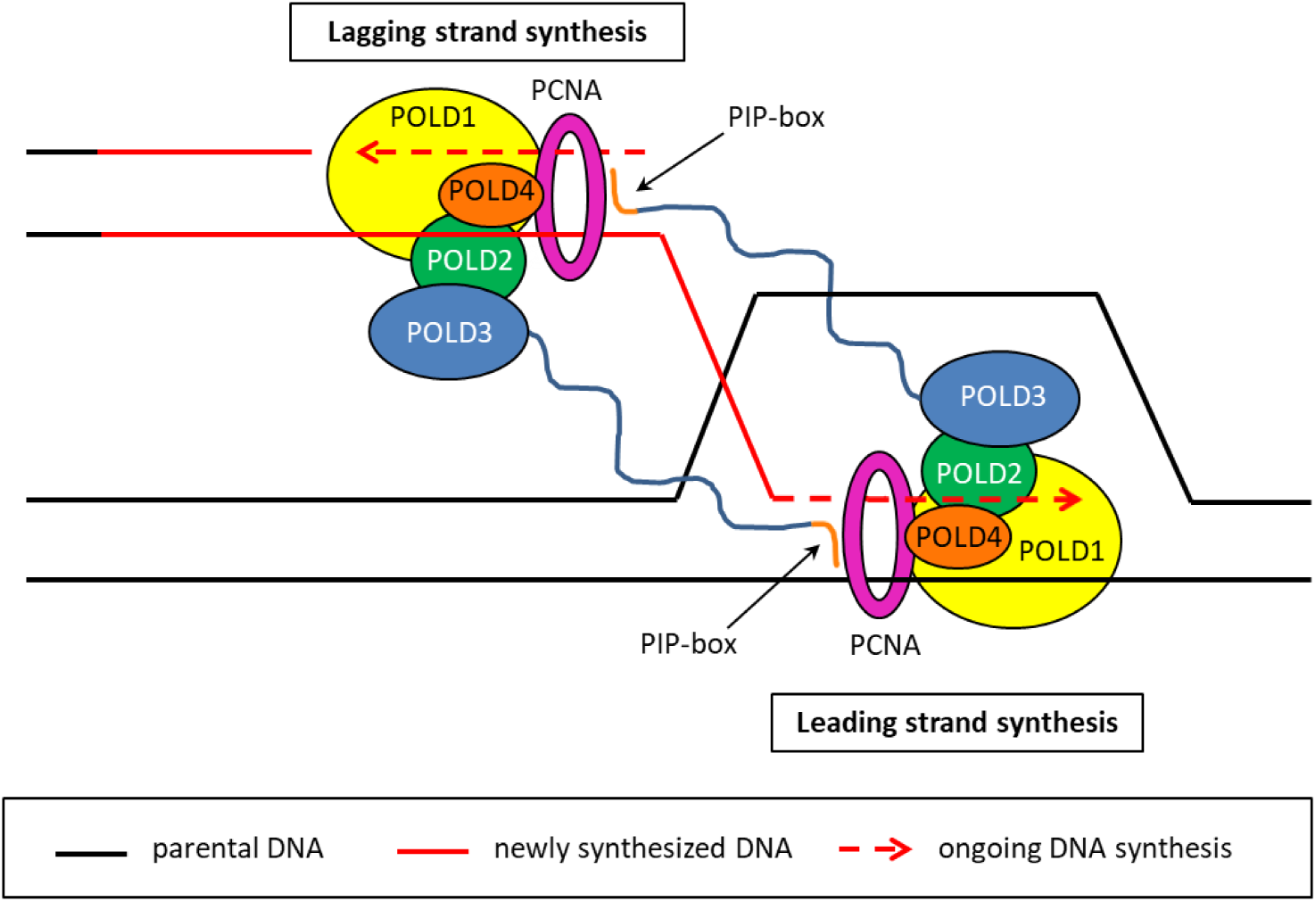
Model proposed on the basis of the results presented in this work. Recent literature suggests that, in MiDAS, DNA polymerase δ might be involved both in leading and lagging strand synthesis, and that the POLD3/PCNA interaction is not involved in the recruit-ment of Polδ at MiDAS sites. Here, we propose that the role of POLD3 in MiDAS might be to co-ordinate leading and lagging strand synthesis. In particular, the C-terminal PIP-box of each of the two POLD3 subunits at the MiDAS fork could bind to the PCNA trimer on the opposite DNA strand, thus providing physical connection between the leading and lagging strand replication ma-chineries.

Due to the importance of MiDAS, and of BIR in general, for cancer cells, a deep understanding of this repair mechanism will provide us with a very promising druggable target for novel cancer therapies. In this work, we provided new insights into the MiDAS-specific role of POLD3, increasing our knowledge about the importance of POLD3 for BIR-related mechanisms.

## MATERIALS AND METHODS

### Cell Culture

HeLa wild type and mutant cells were grown at 37 °C in Dulbecco’s modified Eagle’s medium (DMEM; Invitrogen, Cat. No. 11960) supplemented with 1U/mL penicillin, 1μg/mL streptomycin, 2.92µg/mL glutamine (Invitrogen, Cat. No. 10378-016) and 10% fetal bovine serum (FBS; Invitrogen, Cat. No. 10500).

### CRISPR-Cas9 mutagenesis

For clones 2C4, 2E4, 3H2 and 3D1, guide RNAs were designed manually, synthesized by Integrated DNA Technology™, and resuspended in Nuclease-Free Duplex Buffer (Integrated DNA Technology, Cat. No. 1072570) to get a 100μM solution. To assemble the Cas9:crRNA:tracrRNA ribonucleoprotein complex, 98μL of Nuclease-Free Duplex Buffer were mixed with 1μL of 100μM guide RNA solution and 1μL of a 100μM solution of Alt-R® CRISPR-Cas9 tracrRNA 5’ATTO^TM^ 550 (Integrated DNA Technology, Cat. No. 1077024) in Nuclease-Free Duplex Buffer and warmed up to 95°C for 5 minutes. 66μL of the solution were then mixed with 66μL of 1μM Alt-R™ S.p. HiFi Cas9 Nuclease V3 (Integrated DNA Technology, Cat. No. 1081061) solution in PBS and 68 μL of DMEM. After 10 minutes incubation, the ribonucleotide complex was transfected using the jetCRISPR™ transfection reagent (Polyplus-transfection, Cat. No. 502-07) following the protocol provided by the manufacturer. After 24 hours, cells were collected by trypsinization, and Atto550-positive cells were sorted by MoFlo Astrios Cell Sorter (Beckman Coulter) at the Flow Cytometry facility of the University of Geneva.

As far as clone23 is concerned, a dsDNA oligonucleotide with the sequence coding for the guide RNA was cloned into the pX458 plasmid (Addgene, Cat. No. 48138). After cloning, the plasmid was transfected into HeLa cells using jetPRIME® (Polyplus-transfection, Cat. No. 114-15) as transfection reagent, and GFP-positive cells were sorted by MoFlo Astrios Cell Sorter (Beckman Coulter) at the Flow Cytometry facility of the University of Geneva.

In both cases, single-cell clones obtained after cell sorting were grown in 96-well plates in DMEM complete medium containing 20% FBS until 60-80% confluency, transferred in a 6 well plate in DMEM complete medium containing 10% FBS, and tested for the desired mutation by amplification of the target genomic region by PCR and Sanger sequencing. The sequences of the primers used to test the mutant clones are reported in Supplementary Methods Table 2.

Before proceeding with the production of the mutant clones, the efficiency of each guide RNA was tested with the Alt-R™ Genome Editing Detection Kit (Integrated DNA Technology, Cat. No. 1075932) according to the protocol provided by the manufacturer, using the primers reported in Supplementary Methods Table 1.

### Detection of POLD3 expression in the mutant clones

To check POLD3 expression in the mutant clones, whole cell protein extracts from HeLa wild type and mutant cells were subjected to SDS-PAGE, western blotting and immunoblotting. Cells were collected by trypsinization, washed in PBS, resuspended in 1X EBC buffer (50mM TRIS-HCl pH 8.0, 120mM NaCl, 0.5% NP-40 in water) supplemented with 1mM DTT and 1 pill for 10mL solution of cOmplete™ Protease Inhibitor Cocktail (Roche, Cat. No. 04 693 159 001) and PhosSTOP™ (Roche, Cat. No. 04 906 837 001) and incubated at 4°C for 45 minutes with rotation. Cell lysates were additioned with Benzonase® Nuclease (Millipore, Cat. No. E1014) and 1.5mM MgCl_2_ and incubated at 4°C for 1 hour. Cellular debris were removed by centrifugation, and total protein concentration in the supernatant was measured by Bradford assay (Quick Start™ Bradford 1x Dye Reagent, Bio-Rad, Cat. No. 5000205). Samples were additioned with SDS-PAGE loading dye (final concentration in the sample: 50mM Tris-HCl pH 6.8, 100mM DTT, 2% SDS, 0.1% bromophenol blue and 10% glycerol), warmed up to 95°C for 4 minutes and stored at −20°C. Whole cell extracts were then subjected to SDS-PAGE, and proteins were transferred on a PVDF membrane (Immobilon®-P PVDF Membrane, Millipore, Cat. No. IPVH00010). After over-night blocking at 4°C in Tris Buffered Saline (TBS) supplemented with 0.1% Tween® 20 (PanReac, AppliChem, Cat. No. A4974,0100) and 5% milk, the membrane was incubated with primary antibodies for POLD3 (rabbit, Bethyl, Cat. No. IHC-00249-T, 1:1000 in TBS-0.1% Tween® 20-5% milk) and PCNA (mouse, Genetex, Cat. No. GTX-20029, 1:5000 in TBS-0.1% Tween® 20-5% milk) for 1 hour at room temperature, washed three times in TBS-0.1% Tween® 20, and incubated with HRP conjugate secondary antibodies anti-rabbit (Anti-Rabbit IgG HRP Conjugate, Promega, Cat. No. W4011) and anti-mouse (Anti-Mouse IgG HRP Conjugate, Promega, Cat. No. W4021) diluted 1:2500 in TBS-0.1% Tween® 20-5% milk for 30 minutes. HRP signal was revealed on a chemiluminescence film (Cytiva Amersham™ Hyperfilm™ ECL, GE Healthcare, Cat. No. 28906836).

### MiDAS protocol

The MiDAS experiments were performed by adapting the protocol described in ^22^. Briefly, cells were blocked in G1/S by adding 2mM Thymidine (Sigma-Aldrich, Cat. No. T1895) for 18 hours. After 4 washes in warm PBS, cells were released in S-phase in presence of 0.4 μM Aphidicolin (Cat. No. Sigma-Aldrich, A0781) and of 9 μM RO-3306 (a Cdk1 inhibitor that block cells at the G2/M boundary; Sigma-Aldrich, Cat. No. SML0569) for either 16 hours or 18 hours, the last 1.5 of which in presence of 2 mM HU (Sigma-Aldrich, Cat. No. H8627). After 3 washes in warm media, cells were released in mitosis in presence of 200 ng/mL Nocodazole (Tocris, Cat. No. 1228), 2 mM HU (Sigma-Aldrich, Cat. No. H8627) and 10 μM EdU (Invitrogen, Cat. No. A10044). After either 45, 60 or 90 minutes (in this latter case, EdU was added only for the last 30 minutes), prometaphase and attached cells were collected separately (by mitotic shake-off and trypsinization, respectively) and fixed in 90% methanol over-night for EdU-seq.

### Origin firing and fork progression experiments

To study origin firing and fork progression in the wild type and in the mutant clones, we adapted the protocol described in ^25^. Briefly, cells were blocked in prometaphase by adding 100ng/mL of Noco-dazole to the medium for 8-10 hours. Prometaphase cells were collected by mitotic shake-off and replated in presence of 2mM HU (Sigma-Aldrich, Cat. No. H8627) and 25µM EdU (Invitrogen, Cat. No. A10044) for the origin firing experiments, or only 2mM HU for the fork progression ex-periments. After 13 hours, cells intended for origin firing experiments were collected by trypsiniza-tion and fixed in 90% methanol over-night for EdU-seq. Cells intended for fork progression experi-ments were washed three times with warm PBS and released in media without HU for either 45, 90 or 135 minutes, the last 30 minutes of which in presence of 25µM EdU. Cells were then collected by trypsinization and fixed in 90% methanol over-night for EdU-seq.

### EdU-seq

EdU-labelled DNA isolation and sequencing were performed as described in ^22,25^. Briefly, fixed cells were permeabilized in PBS-0.2% Triton X (PanReac, AppliChem, Cat. No. A1388,2500) and resuspended in click-it reaction mix (85.5mM Tris pH8, 4mM CuSO_4_, 100mM Sodium Ascorbate and 50µM biotin-azide, Jena Biosciences, Cat. No. CLK-A2112-10) for 30 minutes for biotinilation of EdU-labelled DNA. Each sample was then incubated over-night in a solution of 10mM Tris pH 8, 10mM EDTA, 0.5% SDS and proteinase K (Roche, Cat. No. 03115844001) 0.2mg/mL. The next day, proteins and other contaminants were eliminated by phenol-chlorophorm extraction, and puri-fied DNA was precipitated over night at −20°C by adding 0.2M NaCl and two volumes of ethanol to the solution. DNA pellet was washed in 70% ethanol, resuspended in 1xTE (10mM Tris pH 8, 1mM EDTA) and quantified by Qubit ® dsDNA BR assay kit (Invitrogen, Cat. No. Q32853) using a Qubit ® 2.0 Fluorometer (Invitrogen, Cat. No. Q32866). 10μg of DNA from each samples were sonicated by Bioruptor ® Pico sonication device (Diagenode, cat. no. B01060003) to get 100-500bp fragments (6-8 cycles, 15 seconds on, 90 seconds off). EdU-biotin-labelled DNA was isolated by using Dynabeads™ MyOne™ Streptavidin C1 (Invitrogen, Cat. No. 65002) and eluted in Tris 10mM pH8 additioned with 2% β-mercaptoethanol. DNA fragments were then sent to the Ge-nomics Facility of the University of Geneva for library preparation by TruSeq ChIP Sample Prep Kit (Illumina, Cat. No. IP-202-1012) and single-end sequencing by Illumina Hi-Seq 4000 se-quencer.

Sequencing reads were processed and analyzed as in ^25^. Briefly, reads with high quality score were aligned on the human genome assembly (GRCh37/hg19) using the Burrows-Wheeler Aligner (BWA-MEM) software. The aligned reads were assigned to 10kb genomic bins and the sigma val-ues per bin were calculated by using Perl scripts elaborated previously in our group ^25^.

### Calculation of the Index of Directionality (IoD)

To calculate the Index of Directionality (IoD) of each MiDAS peak, the sigma value for forward and reverse reads was extracted for each 10kb bin of each MiDAS region reported for HeLa cells in ^21^. Then, the average sigma value per bin for forward and reverse reads was calculated separately for the right and left parts of the MiDAS region, without taking into account the central 5 bins (the bin reported in ^21^ as MiDAS site center plus the two adjacent bins on its right and left side), the 5 bins on the left and on right end of each MiDAS region, and all the bins which sigma value was lower than 2 for either the forward or reverse reads. Then, the average sigma value for forward and reverse reads for the left and right half peaks were used to calculate the IoD of each MiDAS peak with the formula:

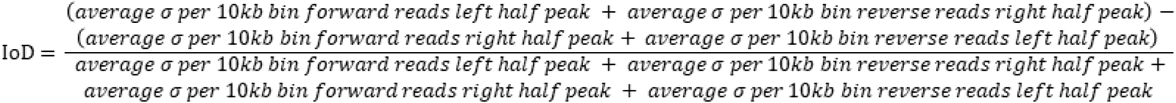

As far as fork progression is concerned, the region used to calculate the IoD spans from the 20 ^th^ bin on the left to the 20^th^ bin on the right of the replication origin center coordinates of U2OS cells pro-vided in ^18^, excluding the central 5 bins and the bins with a sigma value lower than 2.

### Production of average plots for origin firing, fork progression and MiDAS

The average plots for the origin firing and fork progression experiments were produced as in ^18^. Briefly, by using the sigma values calculated from our sequencing data for each 10kb bin, the genome-wide average sigma values in the region spanning +/-0.3Mb from the central bin of every replication origin (provided in ^18^, for U2OS cells) were calculated and plotted.

Average plots for MiDAS sites were produced using the same approach, but average peaks for for-ward and reverse reads were visualized separately. However, since every MiDAS site has a differ-ent size, the MiDAS regions, sourced from ^21^, were stretched or shrunken to the median length of MiDAS sites using the scale-region mode of deepTools ^29^. The plotted region spanned the whole MiDAS sites +/-0.1Mb.

### Analysis of S-phase progression by FACS

Mutant and wild type cells were blocked at G1/S boundary by an 18h treatment with 2mM Thymi-dine. Cells were then washed 4 times with warm PBS and released in warm medium in presence of 10 µM EdU for either 3, 6, 9 or 12 hours. Cells were collected by trypsinization, fixed in 90% methanol overnight and stained for EdU content as described in ^22,25^ for flow cytometry controls. Briefly, after permeabilization in PBS-0.2% Triton X, cells were resuspended for 30 minutes in a click-it reaction as in the “EdU-seq” paragraph above, but containing 1.4μg/mL Alexa Fluor 647 azide (Thermo Fisher Scientific, Cat. No. A10277) instead of biotin-azide. After a PBS wash, cells were incubated overnight in PBS with 6.5μL/mL of RNase, DNAse-free solution (Roche, Cat. No. 11119915001) and 26μL/mL of Propidium Iodide (Sigma, Cat. No. 81845). On the next day, flow cytometry profiles of Propidium Iodide (DNA content) and Alexa Fluor 647 (EdU content) were ac-quired using a Gallios 8-color/2-laser flow cytometer (Beckman Coulter).

## DATA AVAILABILITY

The EdU-seq data produced in the experiments presented in this article have been submitted to the GEO database with accession number: GSE244399.

## AUTHORS CONTRIBUTION

Conceptualization and manuscript writing: D.T. and T.D.H.

Experimental work: D.T. and L.T.

Data processing: V.S.D. and D.T.

Analysis of results: D.T., V.S.D. and T.D.H.

Funding acquisition: T.D.H.

## SUPPLEMENTARY FIGURES

**Supplementary Figure 1:**
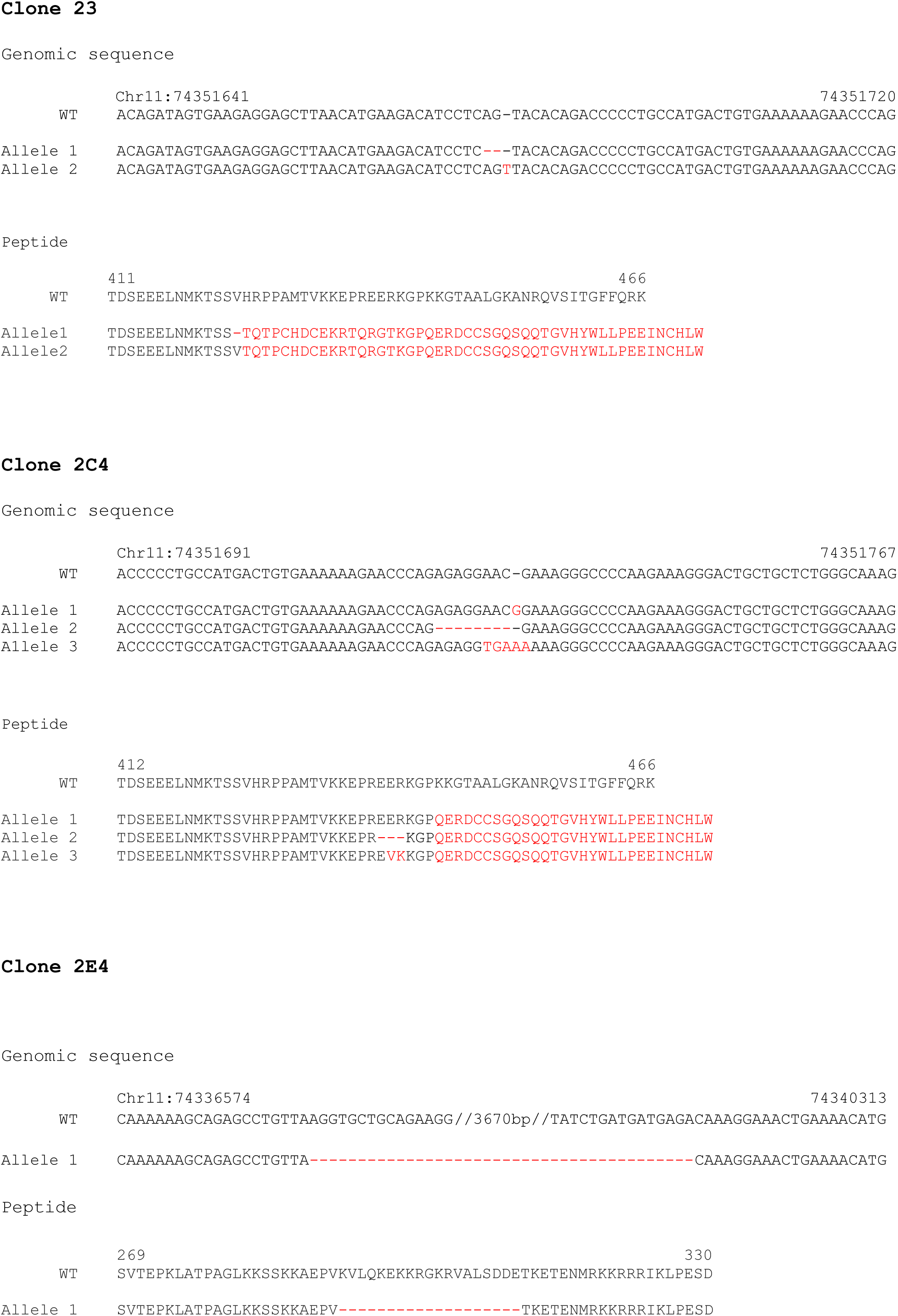
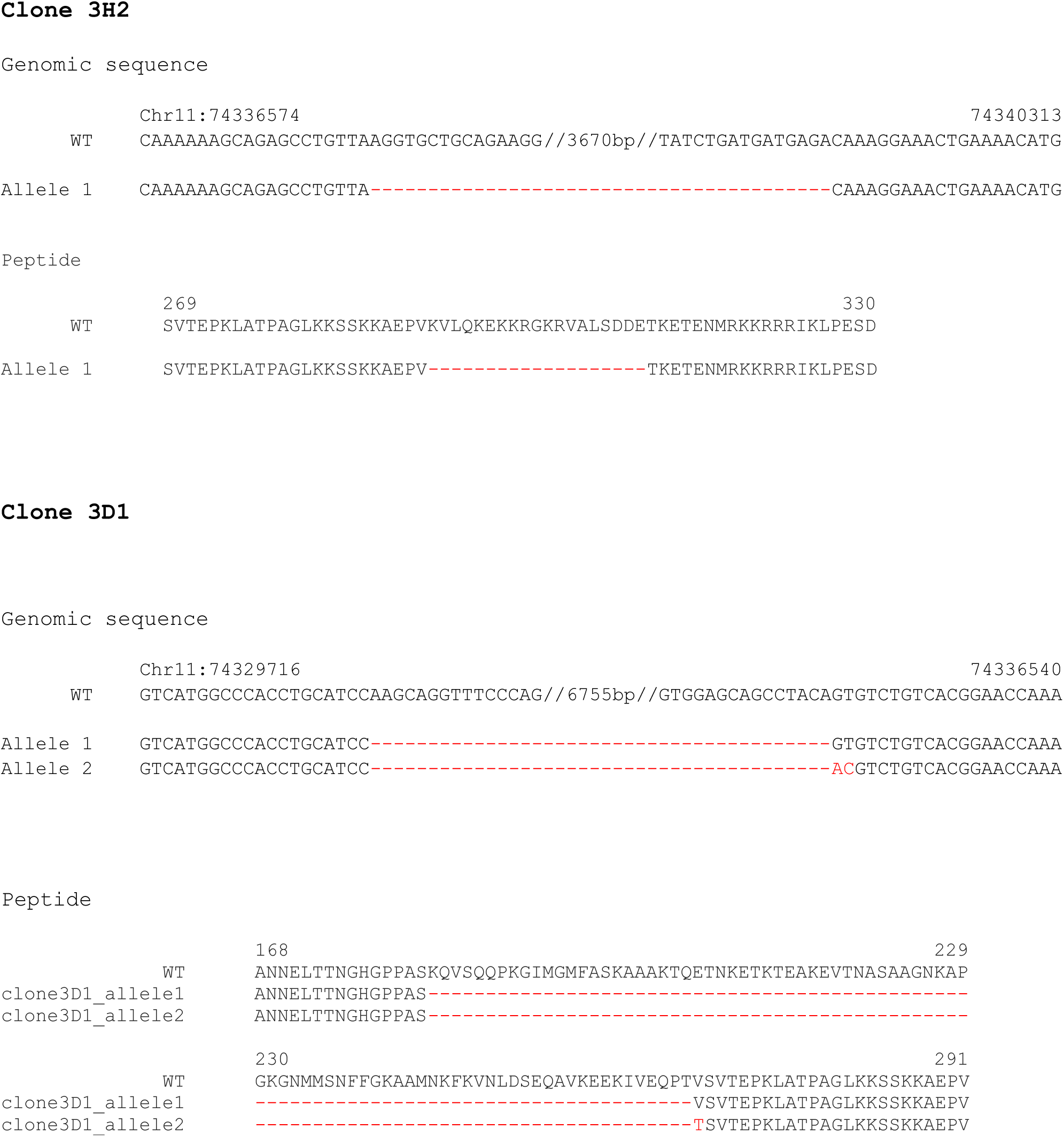
Genomic and protein sequence of POLD3 in the various mutant clones used in this work.

**Supplementary Figure 2:**
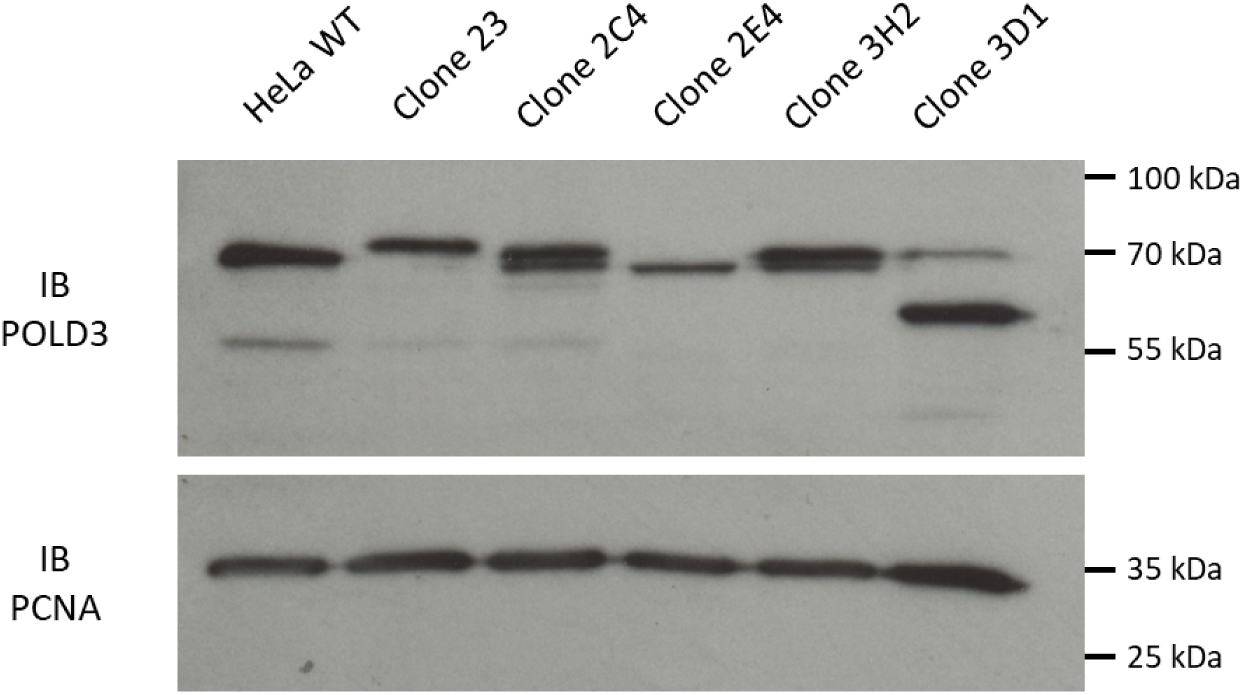
Western Blot for POLD3 and PCNA (as control) performed on whole cell extracts from HeLa wild type and mutant clones. Expected size of the WT protein: 51 kDa; clone 23: 52 kDa; clone 2C4: 52 kDa; clone 2E4: 49 kDa; clone 3H2: 49 kDa; clone 3D1: 42 kDa.

**Supplementary Figure 3:**
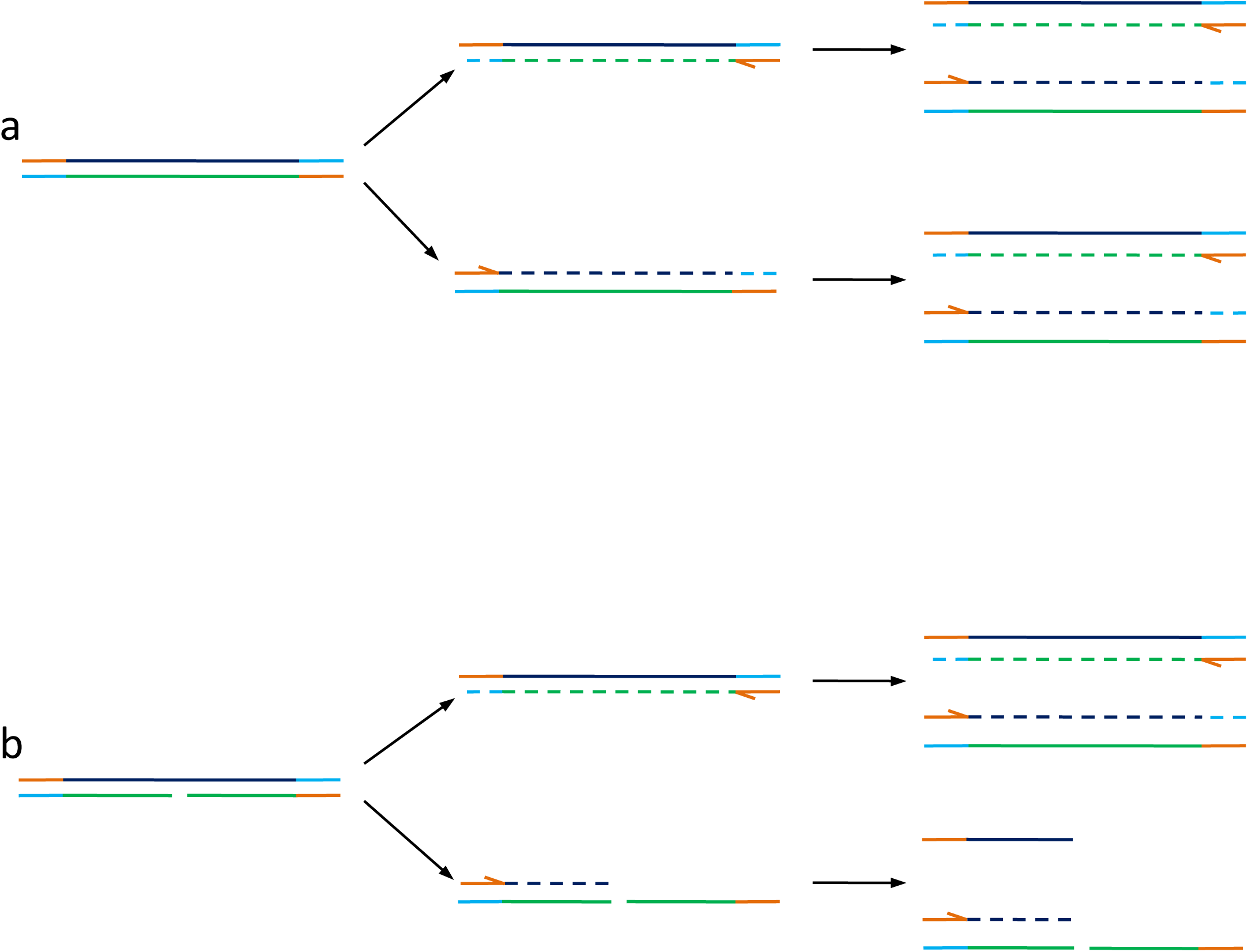
Technical explanation for sequence directionality in the POLD3^delPIP-box^ mutants. **a)** In case of intact dsDNA, both strands are amplified by PCR during library preparation, and ampl-icons coming from both leading and lagging strand are sequenced. **b)** In case of nicked lagging strand, as it might be the case for the POLD3^delPIP-box^ mutants, only the leading strand (intact) is exponentially amplified during library preparation and sequenced, while amplicons coming from the lagging strand (nicked) are poorly represented (they are not exponen-tially amplified) and not even sequenced due to their small dimension (small fragments are ex-cluded from sequencing) and to the lack of one of the two adaptor sequences.

**Supplementary Figure 4:**
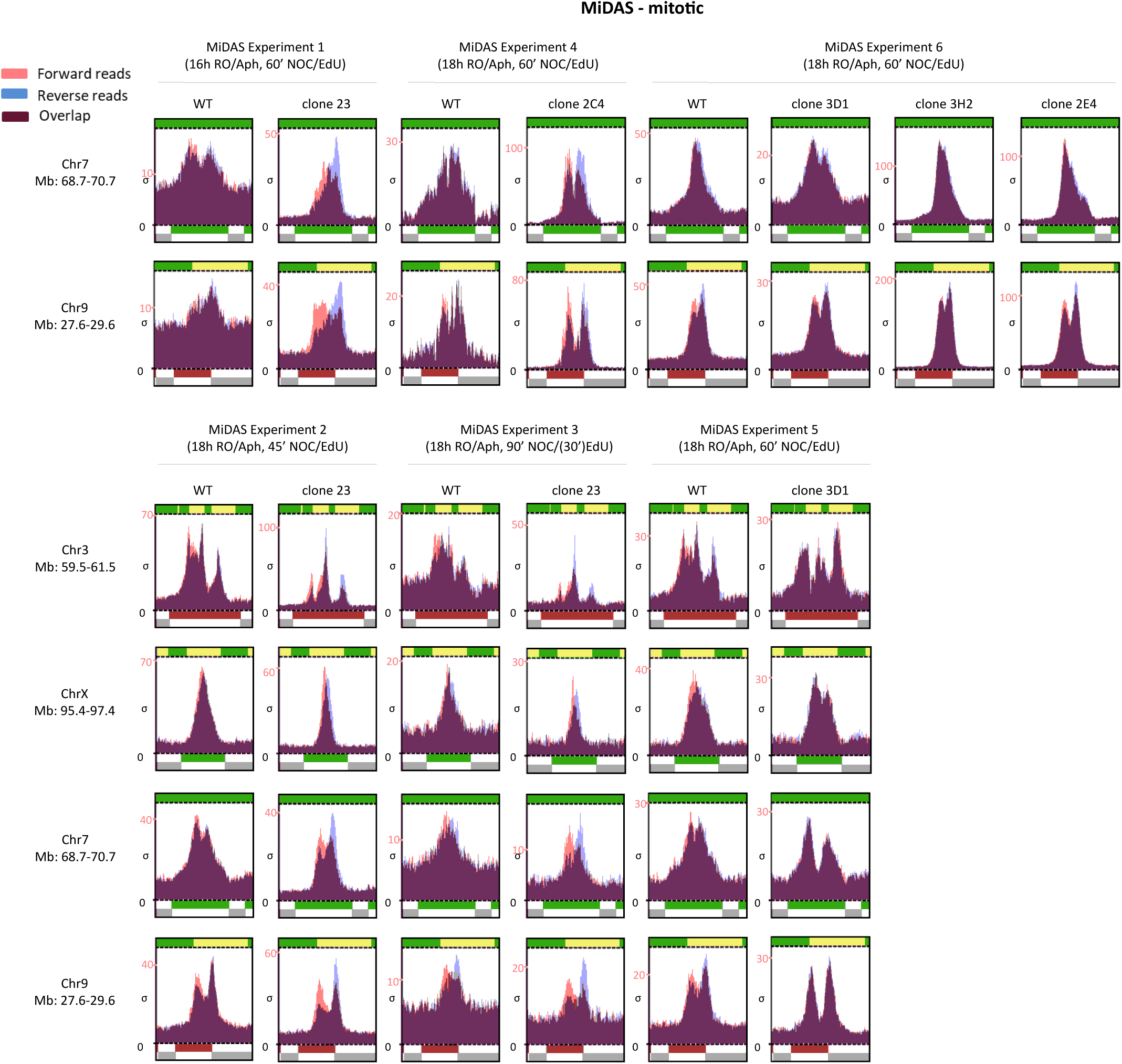
Supplementary results of the MiDAS experiments for mitotic cells. Peaks from forward and reverse reads are shown in red and blue, respectively.

**Supplementary Figure 5:**
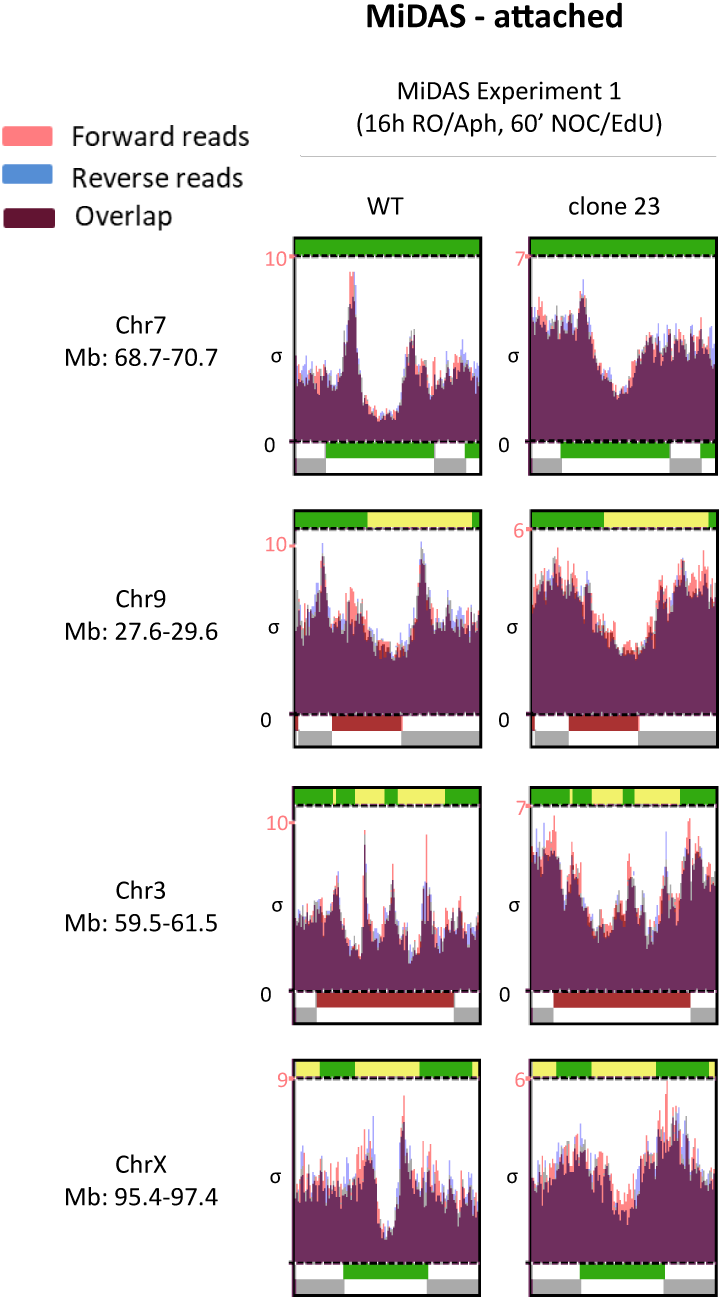
Supplementary results of the MiDAS experiments for attached cells. Peaks from forward and reverse reads are shown in red and blue, respectively.

**Supplementary Figure 6:**
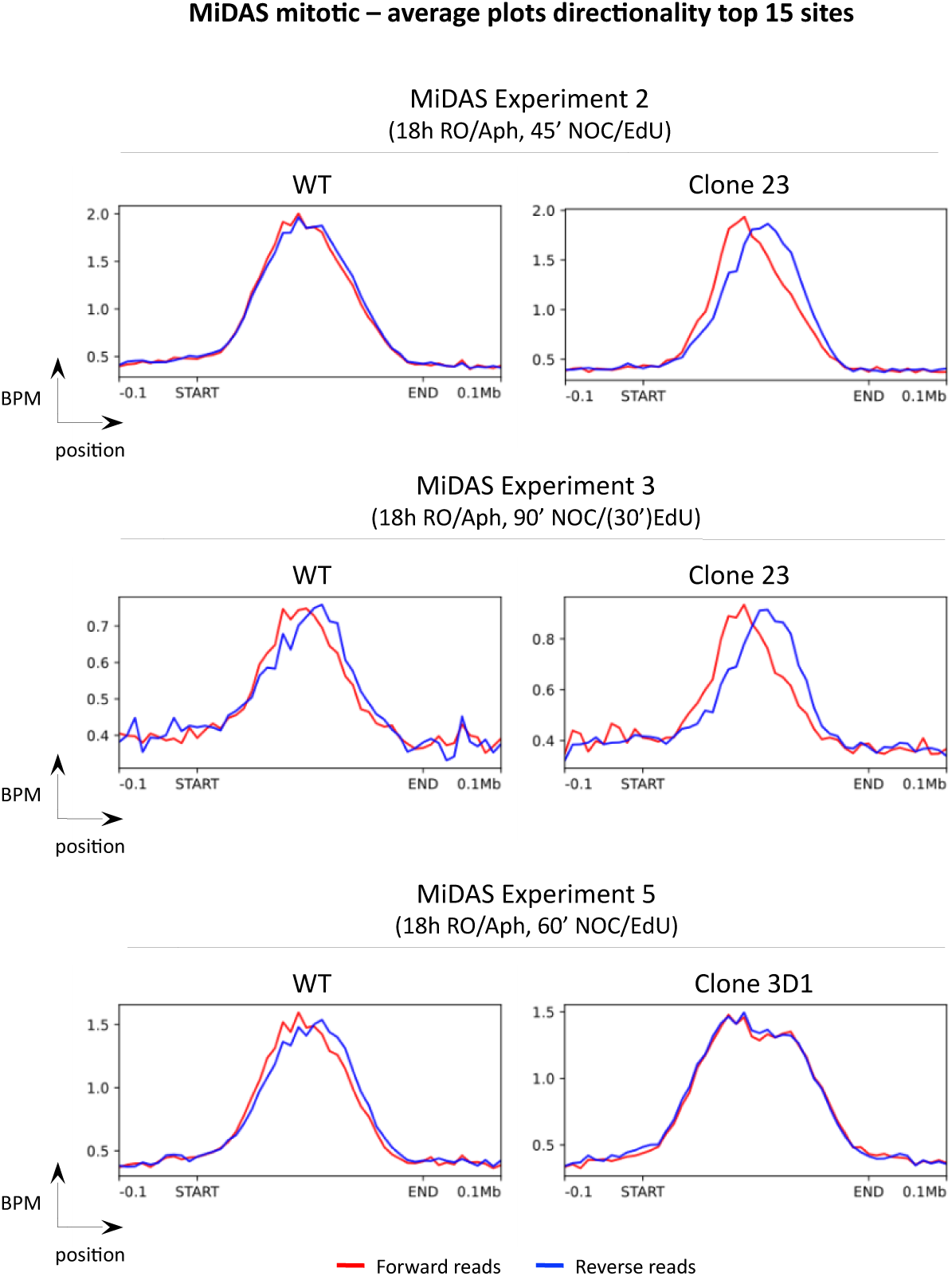
Supplementary average plots of the top 15 MiDAS for the mutant clones and wild type. Average peaks from forward and reverse reads are shown in red and blue, respectively

**Supplementary Figure 7:**
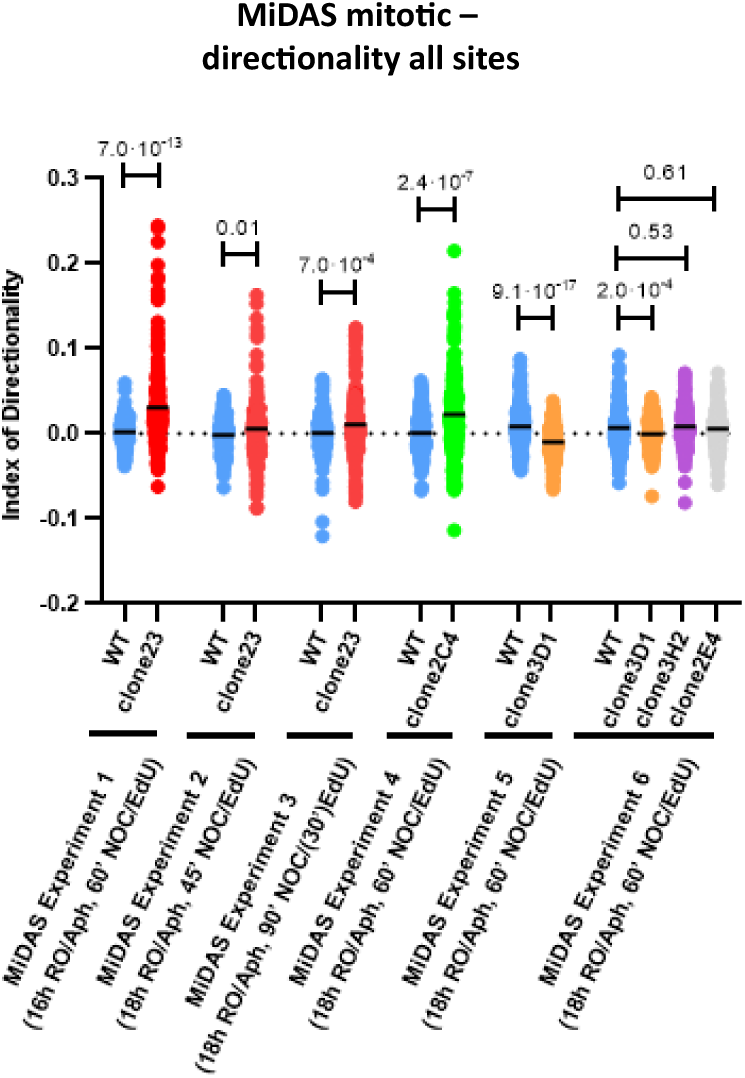
Index of directionality of the MiDAS experiments for the whole set of MiDAS peaks from^21^. The p-values reported on the significance bracket come from a Student’s t-test performed between each mutant clone and the wild type, considering each MiDAS peak as a repeat (statistical significance: p-value<0.05). For each experiment, a horizontal bar indicates the average IoD value of each cell line.

**Supplementary Figure 8:**
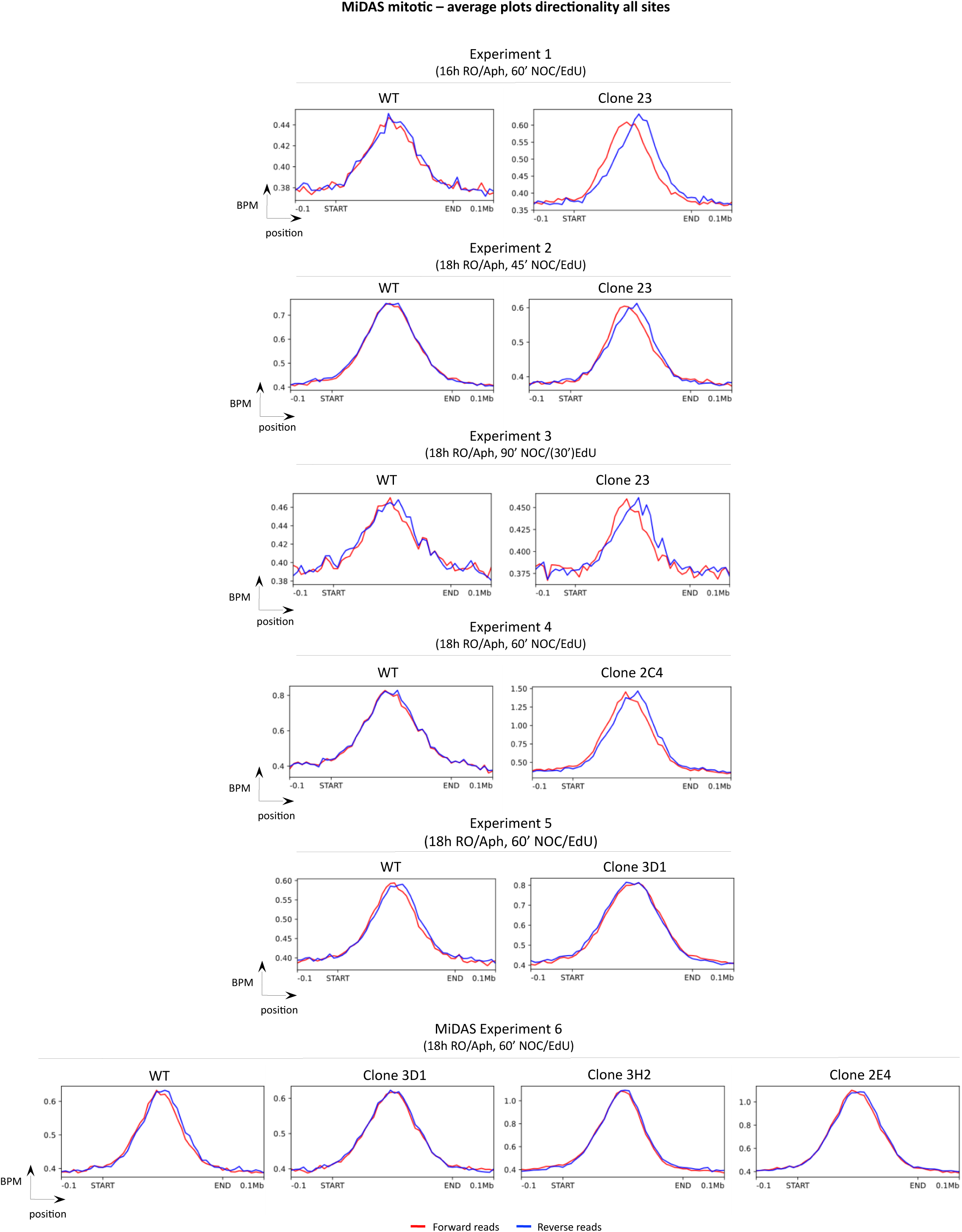
Average plots for the whole set of MiDAS peaks from^21^, for the mutant clones and wild type. Average peaks from forward and reverse reads are shown in red and blue, respectively.

**Supplementary Figure 9:**
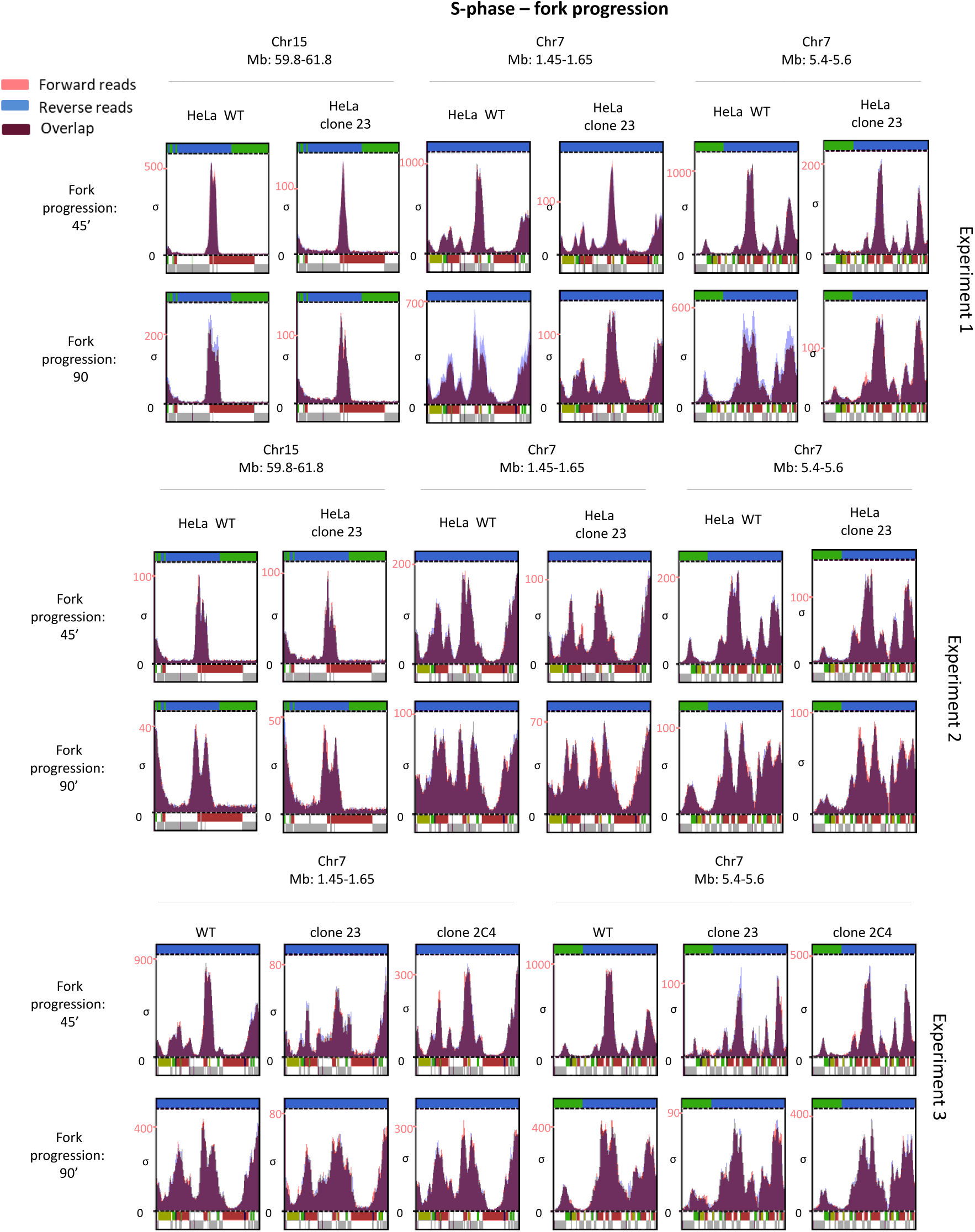
Supplementary results of the origin firing and fork progression experiments. Peaks from forward and reverse reads are shown in red and blue, respectively.

**Supplementary Figure 10:**
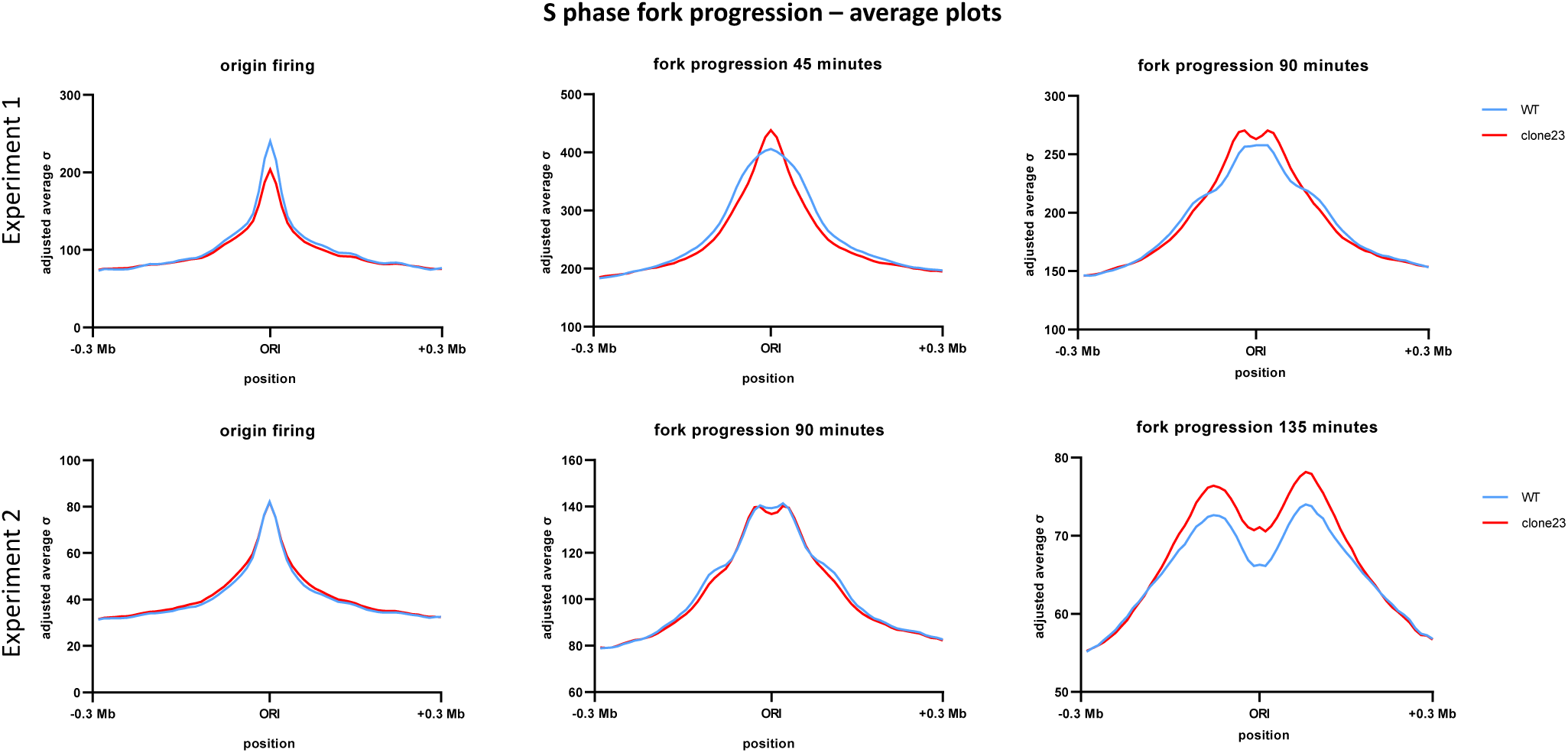
Genome-wide average plots of origin firing and fork progression peaks in clone 23. The average plots have been produced with data from S phase Experiments 1 and 2, using the list of replication origins in U2OS cells from ^18^.

**Supplementary Figure 11:**
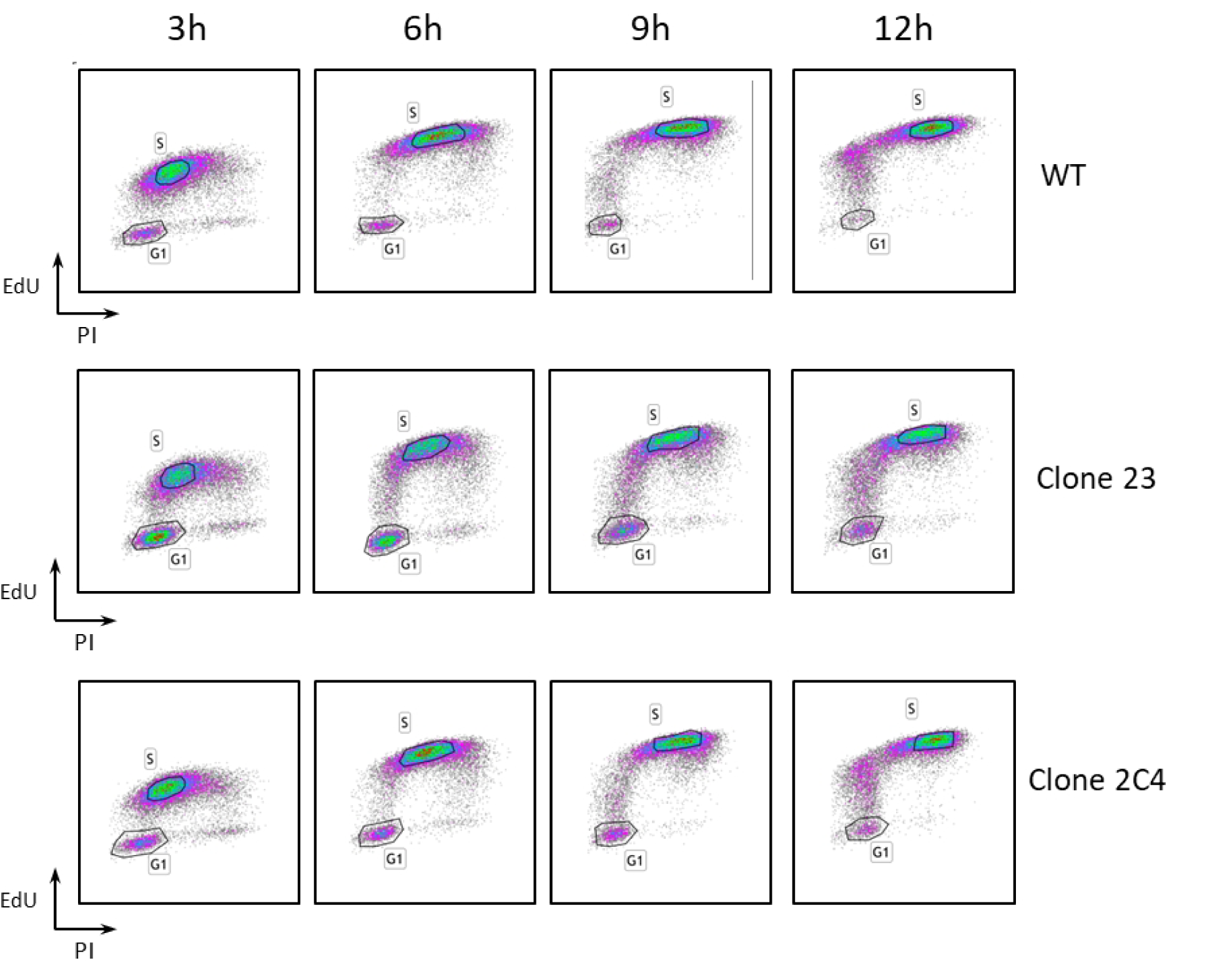
Supplementary results of the FACS experiments performed to investigate S-phase progression in the mutant clones and wild type.

## SUPPLEMENTARY TABLES

**Supplementary Table 1:**
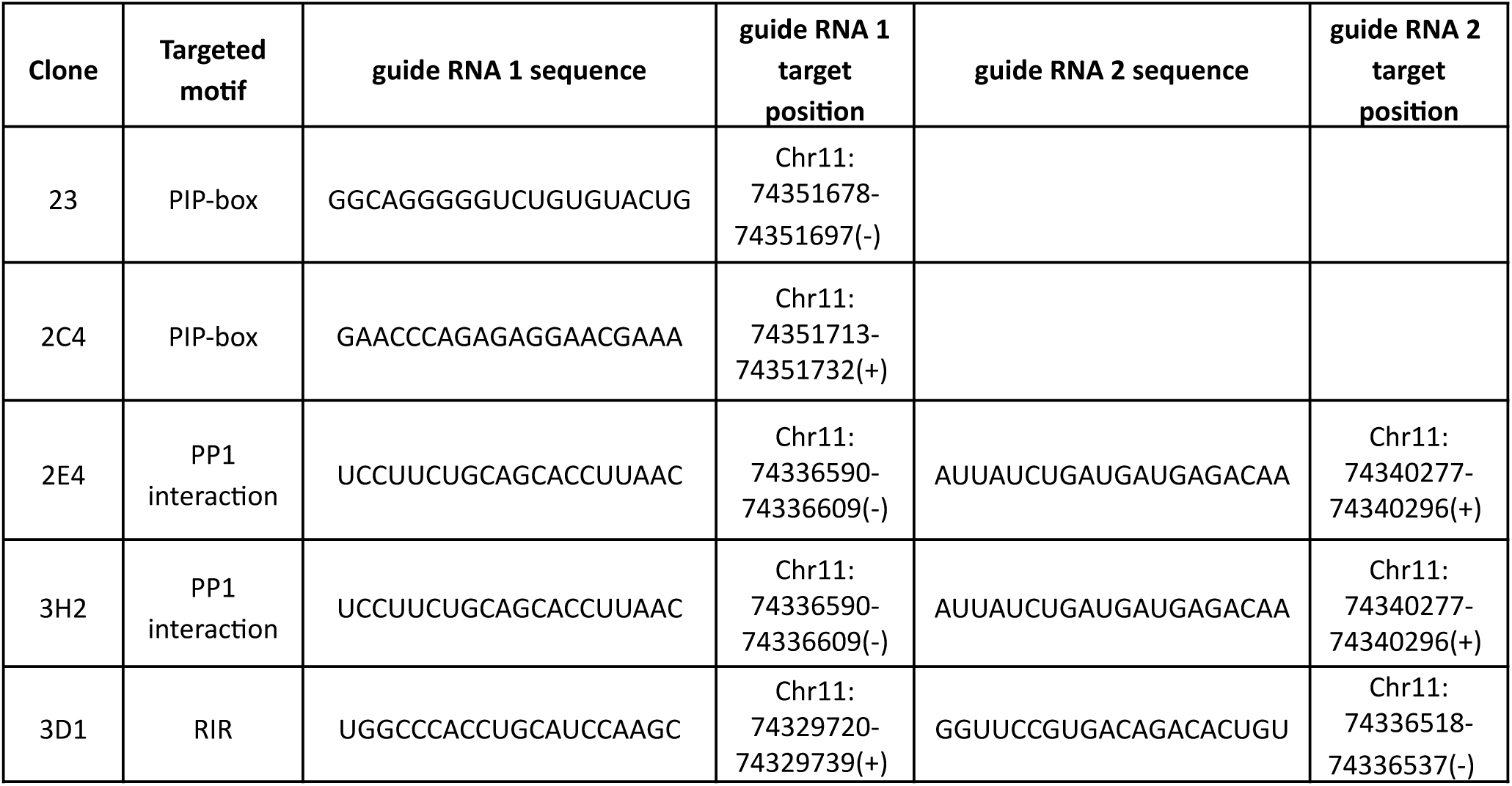
List of the CRISPR guide RNAs used to produce the mutant clones of this work. For each guide RNA used, the sequence and the corresponding genomic positions are provided.

**Supplementary Table 2:** Supplementary statistical analysis for MiDAS directionality in HeLa POLD3^delPIP-box^ clones, based on the top 15 selcted peaks. The average IoD was calculated for the mutant clones and the WT in each MiDAS experiment, and used to perform a paired and an unpaired t test considering each average IoD as a sample.

| Experiment No. | average loD Clone | average loD WT |
| --- | --- | --- |
| MiDAS Experiment 1<br>(Clone 23) | 0.161239193 | 0.02300979 |
| MiDAS Experiment 2<br>(Clone 23) | 0.097779577 | 0.022828017 |
| MiDAS Experiment 3<br>(Clone 23) | 0.089631168 | 0.031475105 |
| MiDAS Experiment 4<br>(Clone 2C4) | 0.13218973 | 0.023051981 |

P-value paired t test: $1.29 \cdot 10^{-2}$
P-value unpaired t test: $1.24 \cdot 10^{-3}$

## SUPPLEMENTARY METHODS TABLES

**Supplementary Methods Table 1:**
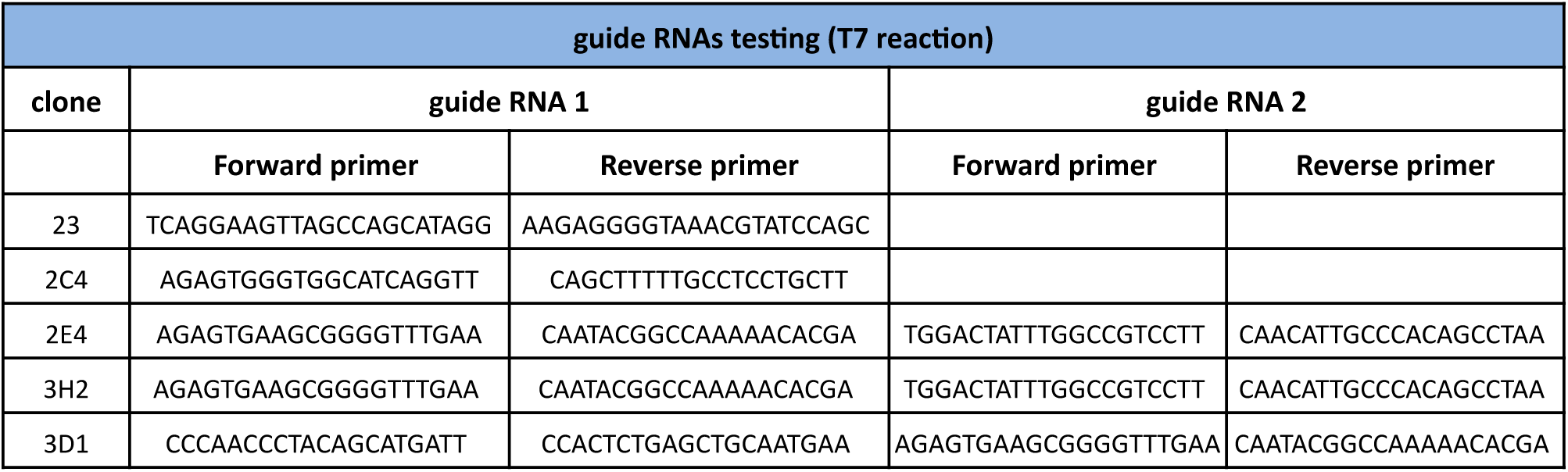
List of the primers used to test the efficiency of the CRISPR guide RNAs by T7 reaction. The sequence and the genomic position of guide RNAs 1 and 2 for each clone are reported in Table S1

**Supplementary Methods Table 2:**
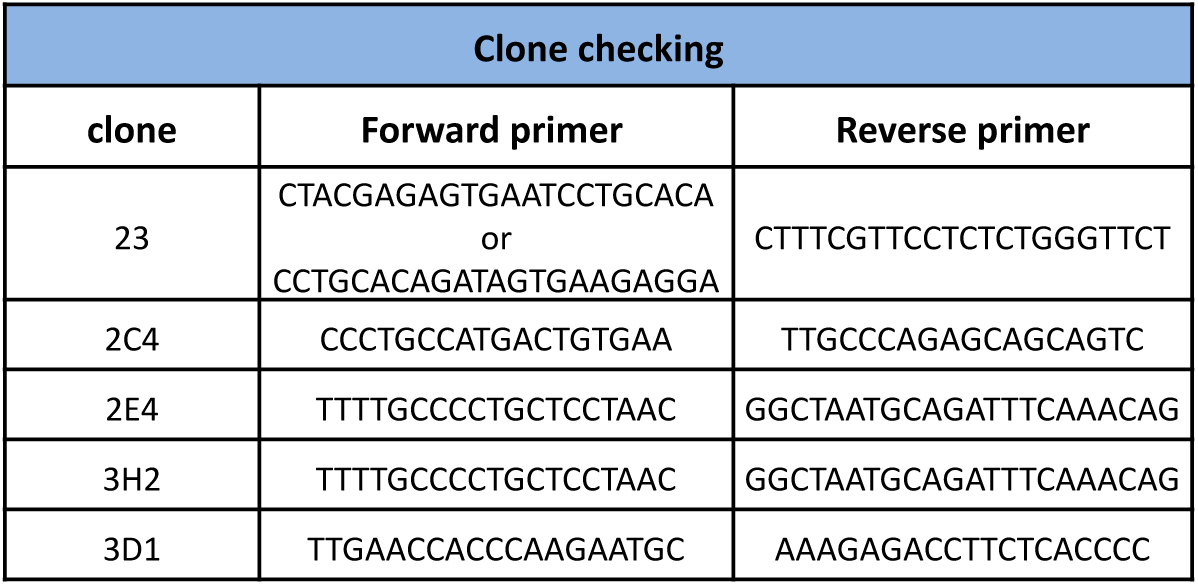
List of the primers used to check the presence of the desired mutations in the mutant clones.

